# PhD and postdoc training outcomes at EMBL: changing career paths for life scientists in Europe

**DOI:** 10.1101/2022.03.01.481975

**Authors:** Junyan Lu, Britta Velten, Bernd Klaus, Mauricio Ramm, Wolfgang Huber, Rachel Coulthard-Graf

## Abstract

The life sciences are training growing numbers of PhDs and postdocs, who increasingly engage in collaborative research. The impact of these changes on the careers of researchers is, however, unclear. Here, we report an analysis of the training outcomes for 2284 researchers who completed a PhD or postdoc at the European Molecular Biology Laboratory (EMBL) between 1997 and 2020. This is the first such study published from a European institute and first time-resolved analysis globally. The most common career outcomes were in academic research, service and teaching (1263 alumni, 55%), including 636 principal investigators (PIs). A broad spectrum of other career paths was also represented, including in industry research (332, 15%) and science-related professions (349, 15%). Our analysis indicates that, although there is increased competition for PI roles, life scientists continue to enter and excel in careers that drive research and innovation.

## Introduction

The career landscape in the life sciences has changed dramatically in the last decades for PhD students and postdocs (collectively referred to in this study as early career researchers (ECRs)). For example, in the US, the percentage of life science doctoral graduates holding a tenure track position 3 to 5 years after graduation more than halved from 18.1% to 8.1% between 1997 and 2015 (1). The principal cause appears to be that the numbers of PhD students and postdocs trained have increased relative to the number of faculty positions (2–6). This trend may stem from economic constraints US institutions faced due to the plateauing of the NIH budget in 2003 after a period of budget increases (7, 8). In Europe, the number of doctorates awarded has also increased faster than number of academic staff, with doctorates increasing by 44% from 82 416 in 2004 to 118 375 in 2012 (9), compared to an increase of only about 9% for full-time equivalent academic staff in the tertiary education sector (10). Economic factors include the stagnation in growth in private, philanthropic and public funding of research and university teaching following the 2008 financial crash (11, 12). Additionally, in both the US and Europe, an increased proportion of funding is now allocated by project-based competitions rather than institutional funding (13–15).

Nevertheless, despite low percentages obtaining these positions, majorities of graduate students starting PhDs have reported aiming for an academic career, with research-intensive principal investigator (PI) roles particularly sought after (16–20). Several surveys have reported high levels of concern from ECRs around career progression, with some linking this to high rates of mental health problems within academia (21–28). Clarity around outcomes is important to help address these concerns and ensure that ECRs can make informed decisions about their career development. Data on training outcomes are also essential for policymakers planning funding and training programs that meet the needs of science, scientists and society as a whole.

In response to calls for increased transparency (29, 30) some North American institutions have published analysis of PhD or postdoc career outcomes (31–33). Via the Coalition for Next Generation Life Science, many additional institutions have also committed to releasing career destinations data and other training outcomes, for example, completion rate and time to degree for PhD students (34). Similarly, in Europe, many universities collect limited information on the initial or current sector of employment (35); a small number of reports have also been released by funders or national agencies, some with analysis of trends in broad sectors of employment (36–38). These reports represent an important step towards transparency, but no detailed time-resolved analysis has yet been published from a university or research institute.

In addition to a changing job market, other changes in research culture may also impact the training of ECRs. For example, there has been an increase in international, interdisciplinary and team-based science (39), with more authors per paper and greater amounts of data included in individual publications (40, 41).

To investigate the effect of these trends on life science training outcomes in Europe, we performed a time-resolved analysis of the career paths of 2284 ECRs who completed a PhD or postdoc at the European Molecular Biology Laboratory (EMBL) between 1997 and 2020. EMBL is an intergovernmental life science research organisation with six sites in Europe. The organisation’s missions include scientific training, basic research in the life sciences, and developing and offering a wide range of scientific services; it currently employs more than 1110 scientists in research and service teams - including over 200 PhD students and 250 postdoctoral fellows focussed on research. EMBL has a long history of training PhD students and postdocs, and was one of the first life science institutes in continental Europe with a structured PhD programme. Fellows in the EMBL International PhD programme have a completion rate of 92% and submit their PhD thesis after 3.95 years on average (data for 2015-2019). More recently, EMBL has launched dedicated fellowship programmes with structured training curricula for postdocs.

This study is, to our knowledge, the first analysis of training outcomes data published by a European research institute. Furthermore, our time-resolved data analysis enables a better understanding of how career outcomes have been changing with time, and how this may extrapolate to the future job market for current ECRs. Finally, our study also links career outcomes to other data, including publication record, to better understand training outcomes.

## Results

Data collection on career and publication outcomes was initially carried out in 2017 and updated in 2021.

Using manual Google searches, we located publicly available information identifying the current role of 89% (2035/2284) of the ECRs in the study (Figure 1A). These alumni were predominantly based in the European Union (60%, 1224/2035), other European countries including UK and Switzerland (20%), and the US (11%). For 71% of alumni (1626/2284), we were able to reconstruct a detailed career path based on online CVs and biographies.

**Figure 1:**
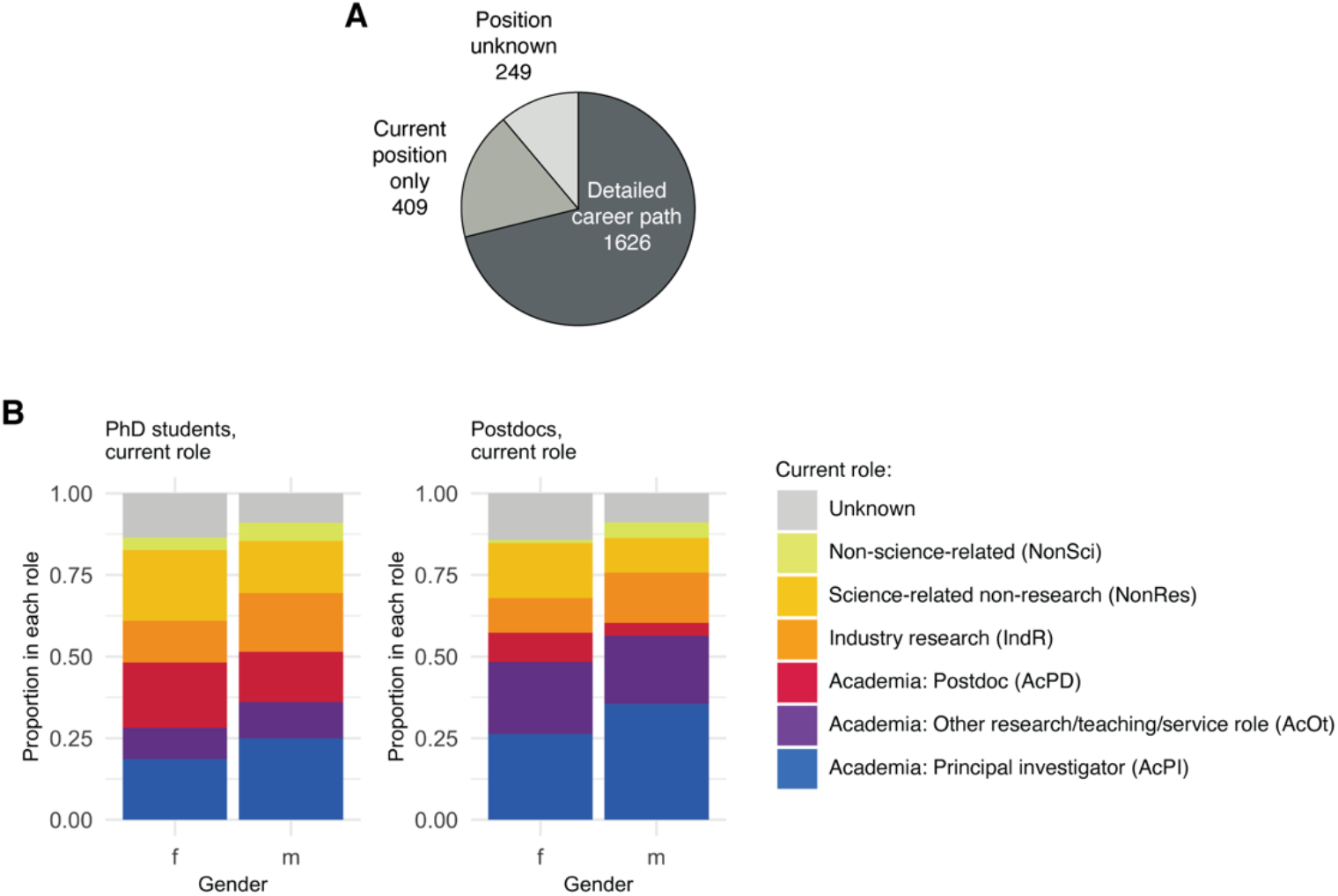
Career outcomes for EMBL PhD and postdoc alumni. **(A)** Pie chart showing data completeness for the 2284 early career researchers (ECRs) included in this study. Detailed career path indicates alumni for whom we were able to reconstruct a career path from EMBL to their current role with a maximum of two 1-year CV gaps, current position only indicates that we were not able to construct a complete career path but were able to identify their current position, and position unknown indicates that we were not able to identify their current role. **(B)** Current role by gender for former PhD students (left, n= 969 [415 female, 554 male]) and postdocs (right, n=1315 [492 female, 823 male]).

On average, ECRs published 4.5 publications from their EMBL work, with 1.6 of those being first author research articles (Supplementary Table 1). Overall, 90% of ECRs (2047/2284) authored at least one publication from their EMBL work, and 73% (1666/2284) published at least one first author research article.

### EMBL alumni contribute to research & innovation in academic and non-academic roles

The majority of alumni (55%) were found to be working in academic research, scientific services and teaching in 2021 (Table 1). An appreciable number of alumni were found employed in industry research (15%) and in science-related roles (15%) not directly carrying out or supervising research, but still closely linked to research, innovation or science education - for example, science communication and patent law. Only 4% were found in professions not closely linked to science. A more detailed summary of the types of role, aligned to a published taxonomy (42), is provided in Supplementary Tables 2-3.

**Table 1:** Overview the current role in 2021 for PhD and postdoc alumni.

| Current role | Role at EMBL |  |  |
| --- | --- | --- | --- |
|  | PhD student | Postdoc | Total |
| Academic research, scientific services & teaching | 485 (50.1%) | 778 (59.2%) | 1263 (55.3%) |
| of which principal Investigator (AcPI) | 215 (22.2%) | 421 (32.0%) | 636 (27.8%) |
| of which other research/service/teaching staff (AcOt) | 102 (10.5%) | 281 (21.4%) | 383 (16.8%) |
| of which postdocs (AcPD) | 168 (17.3%) | 76 (5.8%) | 244 (10.7%) |
| Industry Research (IndR) | 153 (15.8%) | 179 (13.6%) | 332 (14.5%) |
| Science-related non-research professions (NonRes) | 178 (18.4%) | 171 (13.0%) | 349 (15.3%) |
| Non-science-related professions (NonSci) | 47 (4.9%) | 44 (3.3%) | 91 (4.0%) |
| Unknown | 106 (10.9%) | 143 (10.9%) | 249 (10.9%) |
| **Total** | **969** | **1315** | **2284** |

Within academia, 50% of alumni were working as PIs (Table 1), with others working in a variety of roles, including at least 50 alumni in leadership roles within technology platforms and core facilities. The wide variation in job titles between individual companies, sectors and countries impedes the assessment of career progression outside academia. Nevertheless, over 59% of alumni working outside of academic research/service/teaching had a current job title including terminology indicative of a management-level role (manager, leader, senior, head, principal, director, president or chief) (453/766). For leavers from the last five years (2016-2020), this number was 45% (78/174), suggesting that a large proportion of alumni changing sectors enter - or are quickly promoted to - managerial positions.

Within academia, most PI careers follow a linear path: PhD-Postdoc(s)-PI. 95.9% of PIs entered their first PI role from a postdoc (75.3%) or academic research / service / teaching staff role (20.6%), and 93.1% of PIs remained in this type of role (Supplementary Figure 1A, alumni with a detailed career path only). The career paths of those in non-PI roles were less linear. Entry routes for EMBL alumni into industry research positions were, for example, more varied: 20.2% entered their first industry role directly from their PhD, 56.4% from a postdoc position and 13.3% from other non-PI academic positions (Supplementary Figure 1C). Additionally, only 71.6% of those who entered a research position in industry remained in this type of role long-term; 9.9% returned to academia and 18.6% transitioned to non-research or non-science-related professions.

### Gender differences in career outcomes

Many studies have reported that female ECRs are less likely to remain in an academic career, (43, 44). Consistent with these previous studies, we observed differences in PI career outcome by gender for both PhD and postdoc alumni. Female alumni were found less frequently in PI roles and more frequently in non-research science-related positions than their male colleagues (Figure 1B).

### Time-resolved analysis of career outcomes confirms a changing career landscape

For further analysis, alumni were split into three 8-year cohorts. Some demographic differences were seen between cohorts including, at the PhD-level, a higher percentage of female ECRs for more recent cohorts. More recent cohorts were also larger (Table 2).

**Table 2:** Overview of PhD and postdoc cohorts.

| Completion years | PhD cohorts |  |  | Postdoc cohorts |  |  |
| --- | --- | --- | --- | --- | --- | --- |
|  | 1997-2004 | 2005-2012 | 2013-2020 | 1997-2004 | 2005-2012 | 2013-2020 |
| n= | 256 | 341 | 372 | 369 | 422 | 524 |
| % detailed career path | 69.9% | 79.5% | 76.6% | 59.6% | 61.1% | 78.8% |
| % female | 33.2% | 46.0% | 46.5% | 36.9% | 35.3% | 39.5% |

When comparing career outcomes of those with detailed career paths, differences in career outcomes by cohort at equivalent time-points after EMBL are evident (Supplementary Figure 2). For example, the proportion of alumni in PI roles 5-years after EMBL was markedly lower for PhD students and postdocs from the 2005-2012 and 2013-2020 cohorts than the 1997-2004 cohort (Figure 2A). Other smaller changes were also observed in the less well-represented career areas.

**Figure 2:**
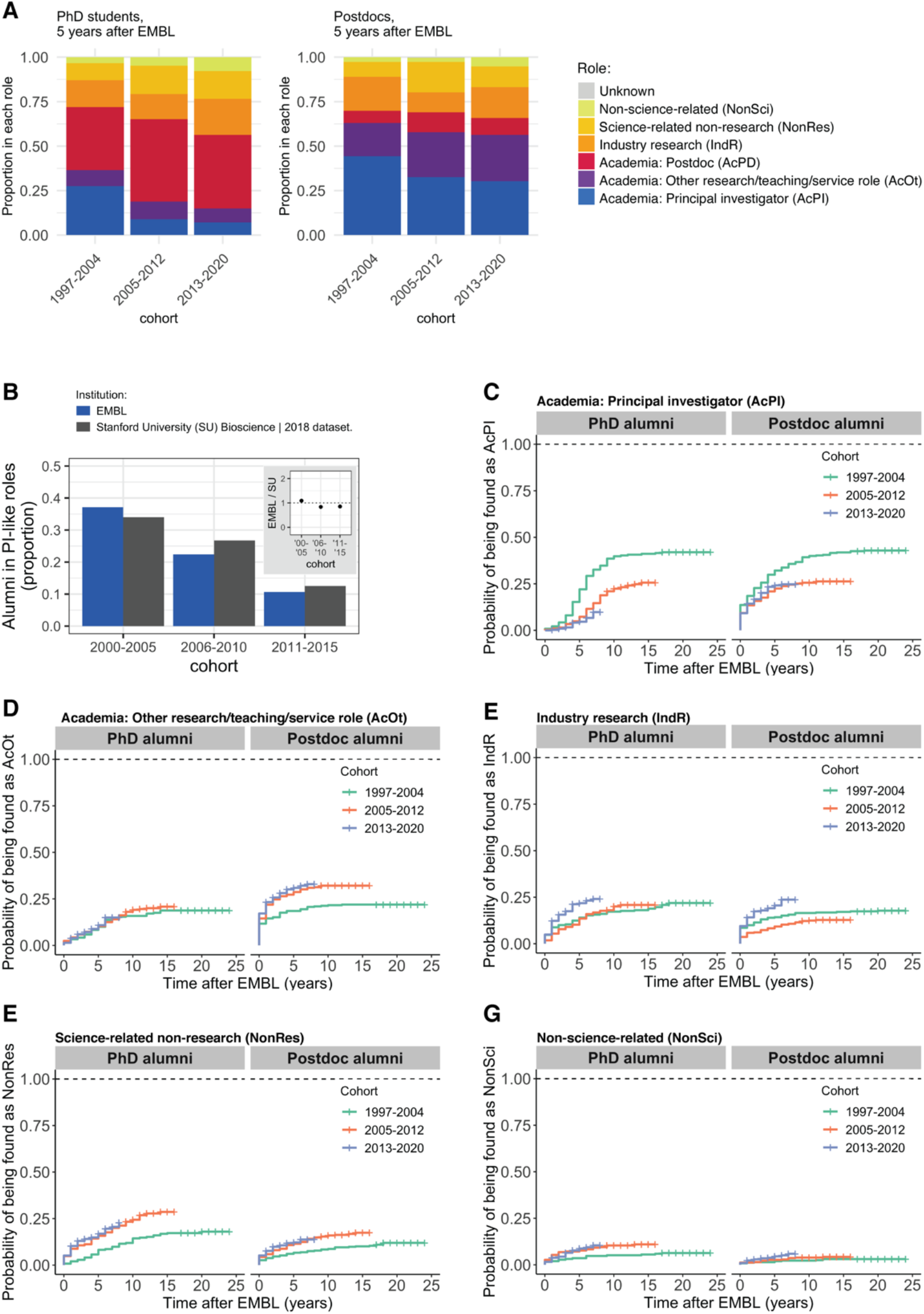
Changes in career outcomes for more recent cohorts. **(A)** Position 5-years after EMBL for PhD students (left, n=617) and postdocs (right, n=686) for whom we have a complete CV and who left EMBL 5-years or more ago, by role classification. **(B)** Column charts showing the proportion of EMBL PhD alumni who were found as principal investigators compared to the publicly available cohort-based data from Stanford University for research-focused faculty (64). Detailed information about the comparison group can be found in Supplementary Table 5. **(C)** Kaplan-Meier plots showing the estimated probability of an individual being found as a PI with time by PhD cohort (left) or Postdoc cohort (right). **(D-G)** as C but for academic research / service / teaching positions (AcOt), Industry Research positions (IndR), non-research science-related positions (NonRes) and non-science-related professions (NonSci).

### The percentage of EMBL alumni becoming PIs is similar to data released by North American institutions for both older and more recent cohorts

To assess whether the changes in career outcomes are EMBL-specific, we compared our data with cohort-based data released by other institutions. We note that there are several limitations in comparing absolute career outcomes between individual studies. Most critically, individual studies often use different data collection methods and role classifications. Furthermore, career outcomes are influenced by the broader scientific ecosystem and the subject focuses of the institution’s departments, which may attract ECRs with dissimilar career motivations. Nevertheless, comparing the outcomes with multiple institutions allows us to interrogate whether the changes we observe for the most frequent, well-defined and linear career path, the PhD-postdoc-PI track, reflect a general trend.

For example, EMBL has strong scientific overlap with Stanford University bioscience departments. Stanford has reported that 34% of its 2000-2005 bioscience PhD alumni and 13% of its 2011-2015 bioscience PhD alumni were in research-focussed faculty roles in 2018(45). The equivalent data for EMBL is similar - with 37% of its 2000-2005 alumni and 11% of its 2011-2015 PhD alumni known to be in PI roles in 2018 (Figure 2B / Supplementary Table 4). Supplementary Figure 3 contains further examples of other programmes with high overlap of scientific areas. Generally, EMBL and the other institutes have a similar proportion of alumni entering PI roles for comparable cohorts. This is consistent with our hypothesis that the differences between cohorts are not EMBL-specific, and reflect a wide-spread change in the number of PhDs and postdocs relative to available PI positions.

We did not analyse the data for non-PI roles, as smaller numbers of individuals entering these roles make it difficult to identify real trends. Additionally, for postdoc career outcomes, only a small number of institutions have released detailed data on the destinations of recent alumni and we are not aware of any long-term cohort-based data.

### While most EMBL ECRs remain in science-related professions, the rate at which they become PIs has decreased with time

To estimate the probability of alumni from different cohorts entering each type of role each year after completing an EMBL PhD or postdoc, we used a statistical regression method, the Cox proportional hazards model. This model is commonly used to model time-to-event distributions from observational data with censoring (i.e., when not all study subjects are monitored until the event occurs, or the event never occurs for some of the subjects). In brief, we fitted the data to a univariate Cox proportional hazards model to calculate hazard ratios, which represent the relative chance of the event considered (here: entering a specific type of role) occurring in each cohort with respect to the oldest cohort. We also calculated Kaplan-Meier estimators, which estimate the probability of the event (entering a specific type of role) at different time points.

For entry of both PhD and postdoc alumni into PI roles, we observe hazard ratios of less than one in the Cox models when comparing the newer cohorts with the oldest cohort (Supplementary Table 5), indicating that the chances of becoming a PI have become lower for the newer cohorts. The Kaplan-Meier curves (Figure 2C) illustrate lower percentages of PIs among alumni from the most recent cohorts compared to the oldest cohort at equivalent time points after completing their EMBL PhD or postdoc. Nevertheless, PI roles remained the most common type of outcome observed for EMBL PhD and postdoc alumni from the 2005-2012 cohort (26.4% and 29.1% found in this career in 2021), and the most recent cohort of alumni appear on a similar trajectory.

For postdoc alumni, our Cox models suggest that an increased proportion of the 2005-2012 and 2013-2020 cohorts entered non-PI academic roles compared to the 1997-2004 cohort (Figure 2D). However, for PhD alumni the rate of entry into non-PI academic roles was similar for all three cohorts.

For both PhD and postdoc alumni, we observed a statistically significant increased probability of entering research roles in industry for the most recent (2013–2020) cohort compared to alumni from 1997-2012 (Figure 2E, Supplementary Table 5). We also saw an increase in numbers entering non-research science-related and non-science-related roles for the 2005-2012 and 2013-2020 cohort compared to 1997-2004 (Figure 2F-G), although this proportion entering non-science-related roles remained small in comparison to other types of roles.

### A small increase in time between PhD conferral year and first PI position is observed

The increased percentage of ECRs entering non-PI careers may fully explain the change in career destinations we observe. However, an increase in total postdoc length could also contribute to the reduced percentage of PIs we observe when comparing career destinations at specific time points after an ECR’s PhD defence. We therefore asked whether an increased duration between PhD and first PI role was observed in our data. In order to fairly compare alumni from different cohorts, we included only alumni for whom we had a detailed career path, who had defended their PhD at least nine-years ago and who had become a PI within nine years of defending their PhD. The nine-year cut-off was chosen as this was the time interval between the last PhDs in the 2005-2012 cohort and the execution of this study; moreover, for PhD alumni from the oldest cohort (defence years 1997-2004), 92% (89/97) of those who became PIs had started this role within nine years of receiving their PhD, indicating that historically most alumni reached this career step within this time period.

From our PhD alumni, 157 alumni met our inclusion criteria for this analysis. On average, this group established their research group 5.6 calendar years (see Methods) after their PhD. There was a statistically significant change in time between PhD and first PI position of 0.8 calendar years between the 1997-2004 cohort (avg. 5.2 years) and 2005-2012 cohort (avg. 6.1 years) (p=0.015) (Supplementary Figure 4A). For postdoc alumni, whilst no statistically significant changes in the time from EMBL to PI role were observed (Supplementary Figure 4B, mean 2.5 years), there was an increase in PhD to PI length of 0.7 calendar years (5.3 calendar years for 1997-2004, 6.0 years for 2005-2012; Supplementary Figure 4C). This may contribute to the reduced rate at which EMBL ECRs become PIs in the first years after EMBL, but is too small to explain the large differences in career trajectories more than 10-years after EMBL, when the number of alumni becoming PIs begins to plateau.

### Publication factors are highly predictive of early entry into a PI position

Publication metrics have been linked to the likelihood of obtaining (46, 47) and succeeding (48) in a faculty position. In this study, alumni who became a PI had more favourable publication metrics from their EMBL work - for example, they published more papers, and papers that had a higher ‘Category Normalized Citation Impact’ (indicating higher numbers of citations compared to other publications in the same field and year; a CNCI of one indicates performance expected by the average paper in that field and year) (Figure 3A-B, Supplementary Table 6).

**Figure 3:**
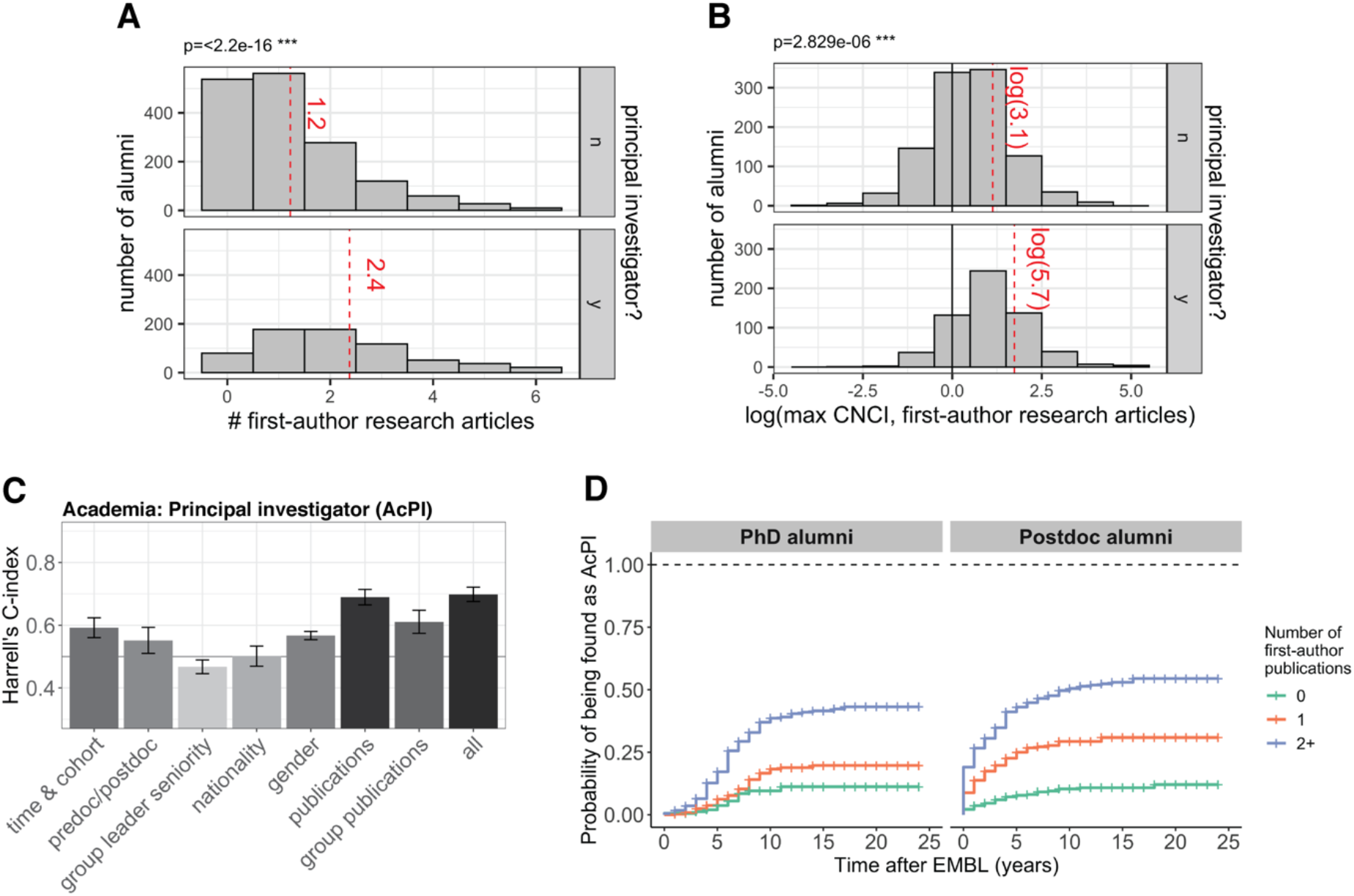
Publication factors are highly correlated with entry into PI positions. **(A)** Histograms of the number of first author EMBL publications per ECR (n=2284) for alumni who have not (yet) become a PI (PI) (top) and those who became a PI (bottom). The number of publications includes only those using an EMBL affiliation. For clearer visualization and to ensure data protection for alumni with outlying numbers of publications, the x-axis (# of first-author research articles) is truncated at the 97.5^th^ percentile. The mean for each group (including outliers) is shown as a red dashed line; the means were compared using a Welch’s t-test. (**B**) Histograms of the natural logarithm of the highest category normalized citation of first author EMBL publications per ECR (n=1656 ECRs publishing at least one first author paper linked to EMBL and with a CNCI value in Clarivate InCites’ database) for alumni who have not (yet) become a PI (top) and those who became a PI (bottom). The log of the mean CNCI for each group is shown as a red dashed line. The distributions were compared using Welch’s t-tests. A CNCI value of 1 (plotted here at log (1)=0)) indicates that the publication’s citations are equal to those expected for the ‘average’ publication in the same research category/categories and year. **(C)** Harrells’ C-Index for models with different sets of covariates for predicting entry into PI positions. A value of 0.5 corresponds to the baseline, i.e., predictive value no better than guessing. A value of above 0.5 indicates that this model has a predictive power, with a value of 1 indicating complete concordance between predicted and observed order to outcome (e.g. entry into a PI position). Values are based on 10-fold cross-validation using a Cox model with ridge penalty. Bars denote the mean across folds and error bars denote the denote the 95% confidence intervals (**D**) Kaplan Meier-curves for probability of becoming PI stratified by number of first-author publications from research completed at EMBL for PhD and postdoc alumni.

To understand the potential contribution of an ECR’s publication record as well as other factors such as cohort, gender, nationality, and seniority of the supervising PI, we fitted multivariate Cox models to the data including these factors as potential predictors for entry into each type of role. We included a range of metrics in our analysis - including journal impact factor, which has been shown to statistically correlate with becoming a PI in some studies (46) and continues to be used by some institutions in research evaluation (49). We, however, wish to note that EMBL is a signatory of the San Francisco Declaration on Research Assessment (DORA), and does not condone the use of journal impact factors for the evaluation of scientists’ work, nor use them in its hiring or evaluation decisions.

To evaluate a model’s predictive power, we used the cross-validated Harrell’s C-index. The C-index measures a model’s predictive power as the average agreement across all pairs of individuals between observed and predicted temporal order of the outcome (for example, becoming a PI). A C-index of 1 indicates complete concordance between observed and predicted order, and 0.5 is the baseline that would be achieved by guessing alone. Prediction is clearly limited by the fact that we could not explicitly encode some covariates that are certain to play an important role in career outcomes, such as career preferences and relevant skills. Nevertheless, the C-index for models containing all data were between 0.61 (entry to Industry Research) and 0.70 (entry into PI roles).

Consistent with previous studies, we found that statistics related to the ECR’s own publications were highly predictive for entry into a PI role: a model containing only the publication statistics performs almost as well as the complete model, reaching a C-index of 0.69 (Figure 3C). Consistent with this, univariate Cox models also suggest that - for example - postdocs with one or more first-author publications from their EMBL work are 3.2 or 6.6 times more likely to become PIs than their colleagues with no first-author publication (Figure 3D, Supplementary Table 7). Group publications (the aggregated publication statistics for all ECRs who were trained within the same research group), cohort/time, gender and type of alumnus (PhD/postdoc), were also predictive for entry into a PI role, with C-indexes of 0.61, 0.59, 0.57 and 0.55, respectively. Models containing only nationality or group leader seniority were not predictive.

Publication statistics for the ECR’s own publications were a weaker predictor of entry into non-PI roles compared to PI roles (Figure 4). For example, for industry research a model containing statistics for the ECRs’ publications had a C-index of 0.54, compared to 0.61 for the complete model (Figure 4B). In univariate Cox models, there were no differences in likelihood of an EMBL PhD with 0, 1 or 2+ publications entering an Industry Research position (4C, Supplementary Table 7). Alumni who published more first author articles were, however, less likely to be found in non-research and non-science roles (Supplementary Figure 5, Supplementary Table 7). Average publication factors for those who enter each type of non-PI role are shown in Supplementary Tables 8-11.

**Figure 4:**
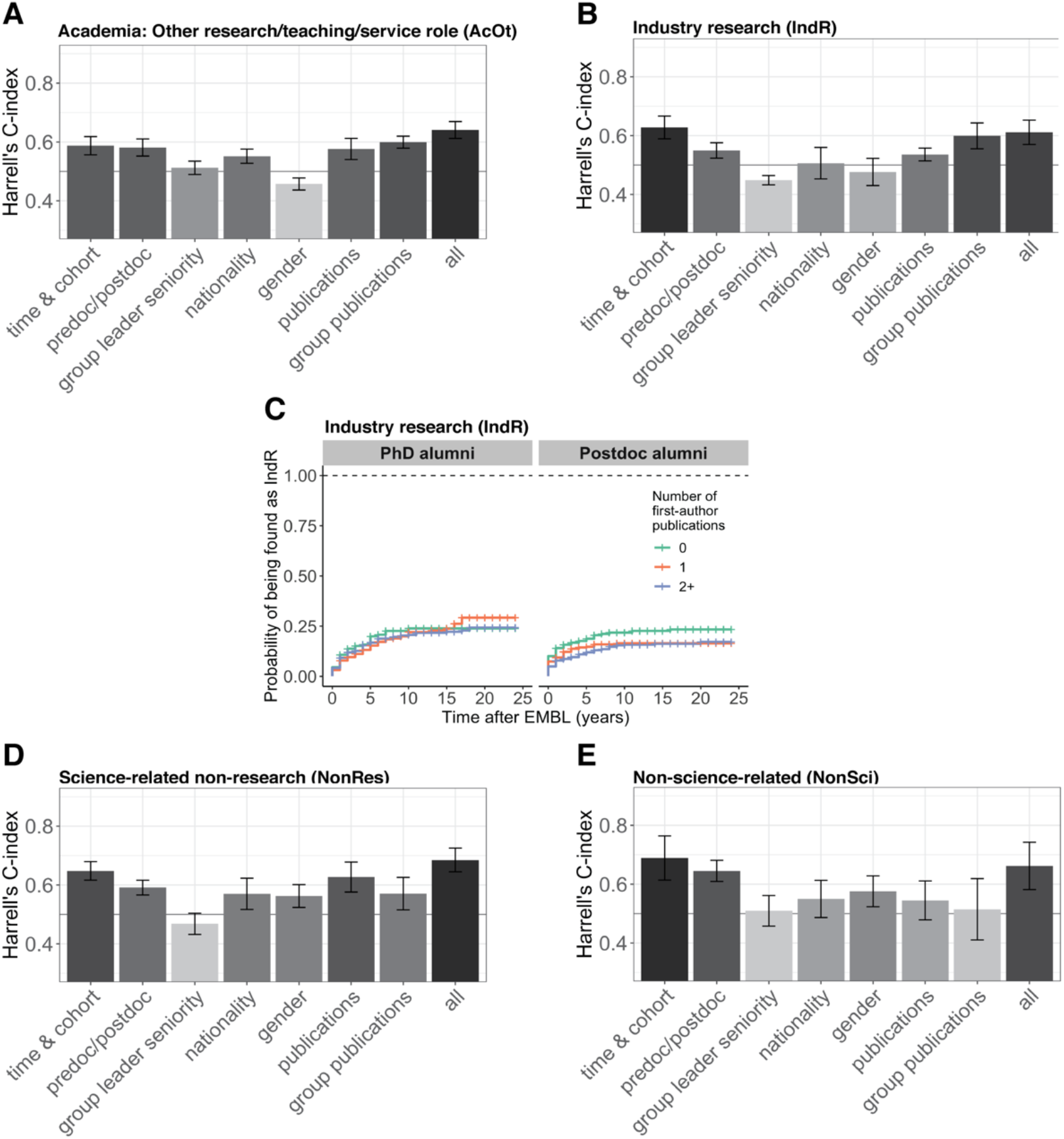
Time-related factors and group publications are most predictive for non-academic career areas and AcOt positions respectively. **(A-B)** Harrells’ C-Index for models sets with different sets of factors for entry into academic research / service / teaching positions (AcOt) (A) and Industry Research positions (IndR)(B). A value of 0.5 is the baseline model, i.e. pure guessing. A value of above 0.5 indicates that this model has a predictive power, with a value of 1 indicating complete concordance between predicted and observed order to outcome (e.g. entry into a PI position). Values are based on 10-fold cross-validation using a Cox model with ridge penalty. Bars denote the mean across folds and error bars denote the 95% confidence intervals (**C**) Kaplan Meier-curves for probability of having entered an Industry Research (IndR) role stratified by number of first-author publications from research completed at EMBL for PhD and postdoc alumni **(D-E)** As A-B but for non-research science-related positions (NonRes) and non-science-related professions (NonSci).

For academic research, service and teaching positions, the factors that were most predictive were those related to the publications of all ECRs in the research group the fellow was trained in (Figure 4A). It is unclear why this might be, but we speculate that this could reflect publication characteristics specific to certain fields that have a high number of staff positions, or other factors such as the scientific reputation, breadth or collaborative nature of the research group and its supervisor. The group’s publications were also predictive for other research and science-related career areas.

Time-related factors (i.e. cohort, PhD award year and EMBL contract start/ end years) were the strongest prediction factors for non-academic career categories (industry research, non-research science-related and non-science related professions, Figure 4B-D).

### More collaborative publications

Previous reports suggest that more recent biology publications have increased amounts of data and more authors (40, 50); a corresponding decrease in the number of first author publications per PhD student has also been reported. For research articles linked to ECRs in this study, the average number of authors per research article has more than doubled between 1995 and 2020 (Figure 5A). No statistically significant difference was found for the number of research articles per ECR between cohorts (Figure 5B, average 3.6 per ECR); however, there was a difference in the number of first author research articles between cohorts. ECRs from the more recent cohorts published fewer first-author publications (Figure 5C, Supplementary Table 12).

**Figure 5:**
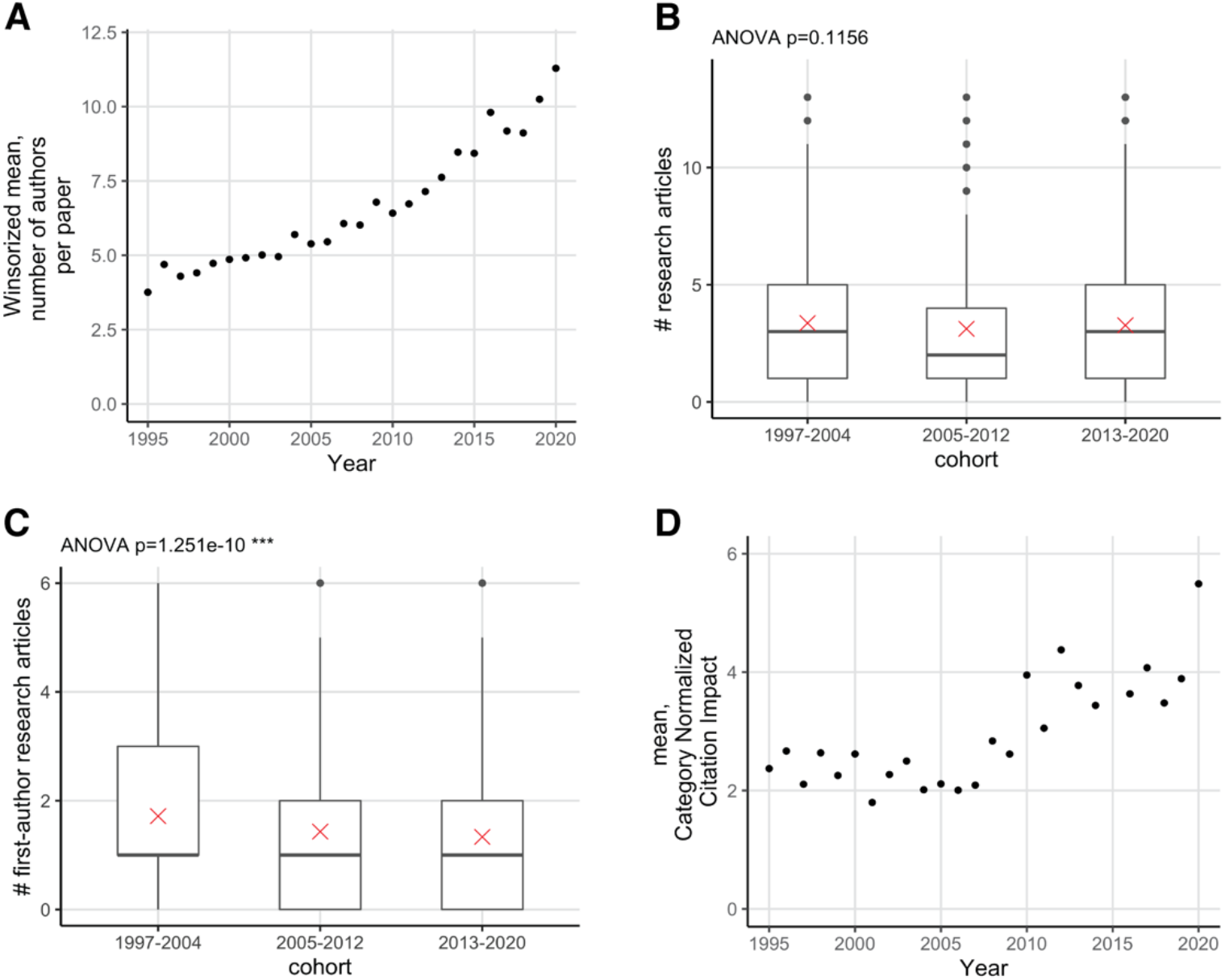
Publications are increasingly collaborative. **(A)** Winsorized mean of authors on EMBL research articles between 1995-2020. Only research articles from 1995-2020 that were assigned to EMBL in the Web of Science InCites database and had at least one author included in this study are shown (n=5413). **(B)** Boxplot showing the distribution of the number of research articles by cohort. The mean is indicated as a red cross. For clearer visualization and to ensure data protection for alumni with outlying numbers of publications, the y-axis of the graph is truncated at the 97.5 percentile. The p-value was generated using one-way analysis of variance (ANOVA) test of the full dataset (including outliers); the p-value excluding outliers is 0.26. **(C)** As B but including first author research articles only; the p-value excluding outliers is 6.5e-7. **(D)** Mean Category Normalized Citation Impact for research articles between 1995-2020 that were assigned to EMBL in the Web of Science InCites database and had at least one author who was included in this study (n=5413).

Data from Clarivate Analytics InCites databases suggest that increasing collaboration may contribute to the increasing number of authors: 79% of EMBL’s publications in 2020 involved international collaboration, compared to 47% in 1995(51). More recent research articles with ECR authors from EMBL (Figure 5D) also had higher ‘Category Normalised Citation Impact’ values.

## Discussion

Our time-resolved analysis of career outcomes indicates that, overall, ECRs in the life sciences have strong job prospects. The career landscape for ECRs has been changing over the last 25 years. Nevertheless, the vast majority of former ECRs continue to pursue careers that contribute to the research and innovation landscape, many in managerial and leadership positions.

Limitations of our study include that its retrospective, observational design limits our ability to disentangle causation from correlation. The changes in career outcomes may, for example, also be influenced by a greater availability or awareness of non-PI career options. EMBL has held an annual career day highlighting non-academic career options since 2006, and many-ECRs actively decide for a career in the private sector, attracted by perceptions of higher pay, more stable contracts, faster impact, and/or better work-life balance. ECRs driven by an interest in specific technologies may also consider roles in research infrastructures and scientific services to be as or more attractive than PI roles. Additionally, we cannot exclude that other factors may also affect the differences we see between cohorts. For example, variations in the number of alumni entering academic roles in countries that offer later scientific independence may have a small effect on outcome data.

To overcome the limitations of this study and better investigate multifactorial and complex issues such as the gender differences in career outcomes, future studies should include large-scale mixed-method longitudinal studies that better record pre-existing career motivations, skills development, and other factors. Multi-institutional studies using consistent data collection methods and a robust classification of roles would also provide a fuller picture of workforce trends.

### Addressing ECR career challenges

ECRs make important contributions to research and develop skills that continue to make them highly employable in academia, industry and other sectors. Nevertheless, some challenges are associated with the diversified career landscape and increase in large-scale projects.

Many ECRs are employed for long time-periods on short-term contracts funded by project-based grants (49) and surveys suggest that ECRs are concerned about career progression (24–28). This may influence high levels of poor mental health amongst ECRs (21, 23). To help alleviate current ECRs’ career concerns and ensure that they can adapt to any further changes in the career landscape - for example, as the result of economic impacts from the corona virus pandemic (52) - it is essential that ECRs are provided with opportunities to reflect on their strengths, understand the wide variety of valuable career options available to them, and develop skills that advance their research projects and employability in their preferred career areas. Efficient and effective support requires input from different stakeholders - including institutions, supervisors, funders, policymakers, and employers of former ECRs - and the engagement of ECRs themselves. At EMBL, a career service was launched in 2019 for all EMBL PhD students and postdocs, building on a successful EC-funded pilot project. In addition to ensuring that career development support is available for all ECRs, policymakers may need to reassess the sustainably of academic career paths, and review the proportion of funding allocated to project-based grants compared to mechanisms that can support PI and non-PI positions with longer-term stability. Providing career development support and more career stability will also support equality, diversity and inclusion in science.

Publication factors are highly predictive of entry into PI careers. For ECRs aiming for a career as a PI, one challenge is balancing the quantity and subjective quality of publications. The observed trend to fewer first author papers likely reflects a global trend towards papers that have more authors, including more international collaborators, as well as EMBL’s focus on ambitious interdisciplinary approaches. Ambitious interdisciplinary projects provide unrivalled opportunities to develop high-performance behaviours including team-work, leadership and creativity. They also allow researchers to tackle challenging biological questions from new angles and deliver publications that significantly advance the field; such publications are seen very positively by academic hiring committees (53–55). However, more complex projects require coordinated input from a large group of co-authors, and may have longer project timelines. Supervisors and ECRs should discuss the potential impact and challenges associated with an ECR’s project, and what can be done to reduce risks.

Research assessment and availability of funding plays an important role in determining funding and career prospects of an academic. Therefore, it is also vital that factors that may affect apparent productivity of ECRs, such as involvement in large-scale projects, career breaks, or time spent on teaching and service, are considered in research assessment. The impact of the coronavirus pandemic on research productivity of researchers with caregiving responsibilities makes these actions imperative (56–58). Initiatives such as the San Francisco Declaration on Research Assessment (DORA) have been advocating for an increased focus on good practice in research assessment, and many funders have reviewed their practices. Cancer Research UK, for example, now asks applicants to its grants to describe three to five research achievements, which can include non-publication outputs (59). Other initiatives include more transparent author contribution information in publications (60, 61) and promotion of “FAIR” principles of data management (62). We also expect the increasing use of preprints (40, 63) to have a positive effect on the careers of ECRs involved in ambitious projects with longer publication time-scales.

## Conclusions

Our data highlight the many ways that early career researchers contribute to the research and innovation landscape, and suggest that early career life scientists continue to enter leadership roles in academia, industry and other sectors. ECR training therefore continues to be a valuable investment that creates a highly qualified workforce with strong job prospects.

Nevertheless, a number of challenges exist for ECRs in their careers, particularly increasing competition for mid-career research leadership roles in academia. These challenges may also increase in the coming years due to the impacts of the coronavirus pandemic. Adequate support and policies to address these challenges are therefore urgently needed. Continued monitoring of career outcomes will be essential for policymakers deciding how best to adapt funding and training programmes to support sustainable career paths in the life sciences, and to enable ECRs to make informed decisions about their own career development.

## Methods

### Data collection & analysis

The study includes individuals who graduated from the EMBL International PhD Programme (n=969) between 1997 and 2020 or who left the EMBL postdoc programme between 1997 and 2020 after spending at least one year as an EMBL postdoctoral fellow (n=1315). Each person is included only once in the study: where a PhD student remained at EMBL for a bridging or longer postdoc, they were included as PhD alumni only, with the postdoc position listed as a career outcome.

For each alumnus or alumna, we retrieved demographic information from our internal records and identified publicly available information about each person’s career path (see supplementary methods). Where possible, this information was used to reconstruct a detailed career path for each ECR. An individual was classified as having a “detailed career path” if an online CV or biosketch was found that accounted for their time since EMBL excluding a maximum of two 1-calendar-year career breaks (which may, for example, reflect undisclosed sabbaticals or parental leave). Each position was classified using a detailed taxonomy, based on a published schema (42), and given a broad overall classification (see Supplementary Methods). The country of the position was also recorded. For the most recent position, we noted whether the job title was indicative of a senior or management level role (included “VP”, “chief’,”cso”,”cto”;”ceo”;”head;”principal”;”president”;”manager”; “leader”; “senior”), or if they appeared to be running a scientific service or core facility in academia.

We use calendar years for all outcome data - for example, for an ECR who left EMBL in 2012, the position one calendar year after EMBL would be the position held in 2013. If multiple positions were held in that year, we take the most recent position. We use calendar years, as the available online information often only provides the start and end year of a position (rather than exact date).

An EMBL publication record was also reconstituted for each person in the study. Each of their publications linked to EMBL in Clarivate Analytics’ Web of Science and InCites databases in June 2021 were recorded. The data included publication year and - for those indexed in InCites-crude metrics, such as category normalised citation impact and percentile in subject area (measures of citations) as well as journal impact factor. EMBL publications were assigned to individuals in the study based on matching name and publication year (see Supplementary Methods for full description). When an individual was the second author on a publication, we manually checked for declarations of co-first authorship. Aggregate publication statistics for ECRs with the same primary supervisor were also calculated.

The names and other demographic information that would allow easy identification of individuals in the case of a data breach were pseudonymised. A file with key data for analysis and visualisation in R was then generated. A description of this data table can be found in Supplementary Table 1, along with summary statistics.

### Statistical model

A Cox proportional hazards regression model was fitted to the data in order to predict time-to-event probabilities for each type of career outcome based on different covariates including cohort, publication variables and gender. Multivariate Cox models were fitted using a ridge penalty with penalty parameter chosen by 10-fold cross-validation. Harrell’s C-index was calculated for each fit in an outer cross-validation scheme for validation and analysis of different models, with 10-fold cross-validation.

### Data availability, and data protection

The data were collated for the provision of statistics, and are stored in a manner compliant with EMBL’s internal policy on data protection (https://www.embl.org/documents/document/internal-policy-no-68-on-general-data-protection/). The nature of the data precludes sufficient anonymisation to provide a public data-release. Summary statistics for the data table used to generate the figures can be found in Supplementary Table 1.

### Code availability

Rmarkdown documentation of the analysis and figures (except 1A, created in adobe illustrator) can be found at: https://www.huber.embl.de/users/jlu/EMBLcareer/analysis.html

Source code is available at: https://github.com/Huber-group-EMBL/EMBL-Career-Analysis

## Acknowledgements

We would like to thank Monika Lachner and Anne Ephrussi for their critical reading of the manuscript and strong support of this project. We also acknowledge the instrumental support of EMBL’s Alumni Relations, HR, SAP, Library, International PhD Programme and Postdoc Programme teams providing the initial information that allowed this study to be completed. Finally, we thank Edith Heard, Brenda Stride, Jana Watson-Kapps (FMI), and EMBL’s Directorate, SAC, SSMAC and Council for discussion.

Rachel Coulthard-Graf is employed by EMBL’s Interdisciplinary Postdoc Programme, which has received funding from the European Union’s Horizon 2020 research and innovation programme under Marie Skłodowska-Curie grant agreements 664726 (2016-2020) and 847543 (2019-present).

## Author contributions

Junyan Lu: Methodology; Formal analysis; Data Curation; Writing - review & editing; Visualization. Britta Velten: Methodology; Formal analysis; Data Curation; Writing - review & editing; Visualization. Bernd Klaus: Methodology; Formal analysis; Data Curation; Writing - review & editing; Visualization. Mauricio Ramm: Methodology; Investigation. Wolfgang Huber: Methodology; Writing - review & editing; Supervision. Rachel Coulthard-Graf: Conceptualization; Methodology; Formal analysis; Investigation; Data curation; Writing - original draft preparation; Writing - review & editing; Visualization.

