## SupplementaryFiguresTablesMethods for "PhD and postdoc training outcomes at EMBL: changing career paths for life scientists in Europe"

#### Supplementary figures, tables and methods

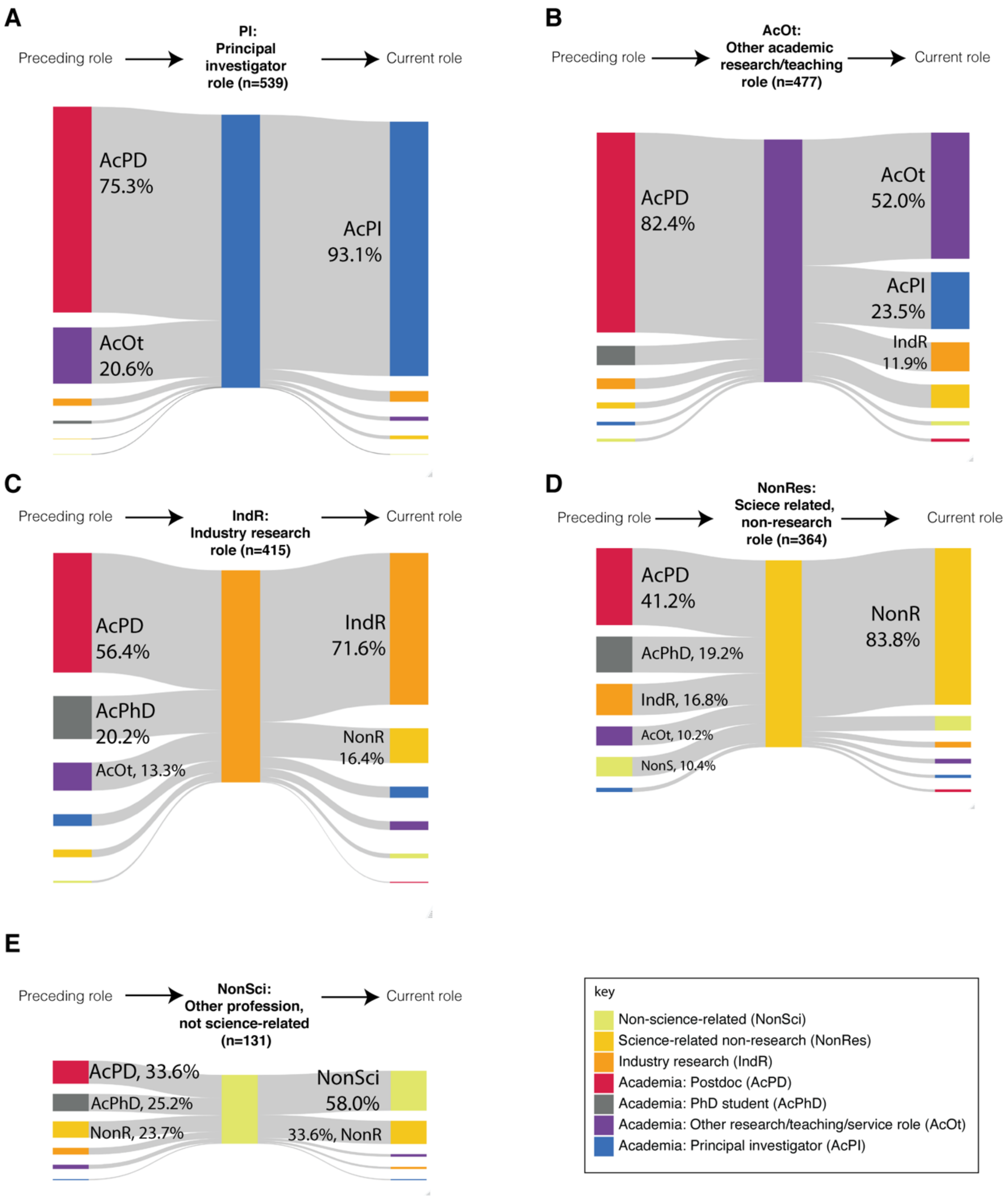

**Supplementary Figure 1: Simplified career paths for alumni who have held different types of role** Sankey diagrams focussing on different types of role, showing the position alumni had before entering their first role of that type (left bar), and what their current role is (right bar). Data is shown only for alumni for whom a detailed career path is available. A preceding AcPD (academic postdoc) role includes both entry directly from an EMBL postdoc and entry via a postdoc position held after leaving EMBL. For EMBL PhD students, a bridging postdoc position in the same lab as the PhD is included within the AcPhD category, rather than AcPD. Category labels with percentages are shown for values of 10% or higher. Diagrams were created in R and scaled manually so that the bar height is proportional to the number of alumni in the role. (A) Principal investigator (PI, blue) n=539 (B) Other academic research / teaching roles (AcOt, purple) n=477 (C) Industry research (IndR, orange) n=415 (D) Science-related, non-research roles (NonRes, yellow) n=364 (E) Non-science-related roles (NonSci, light yellow-green) n=131.

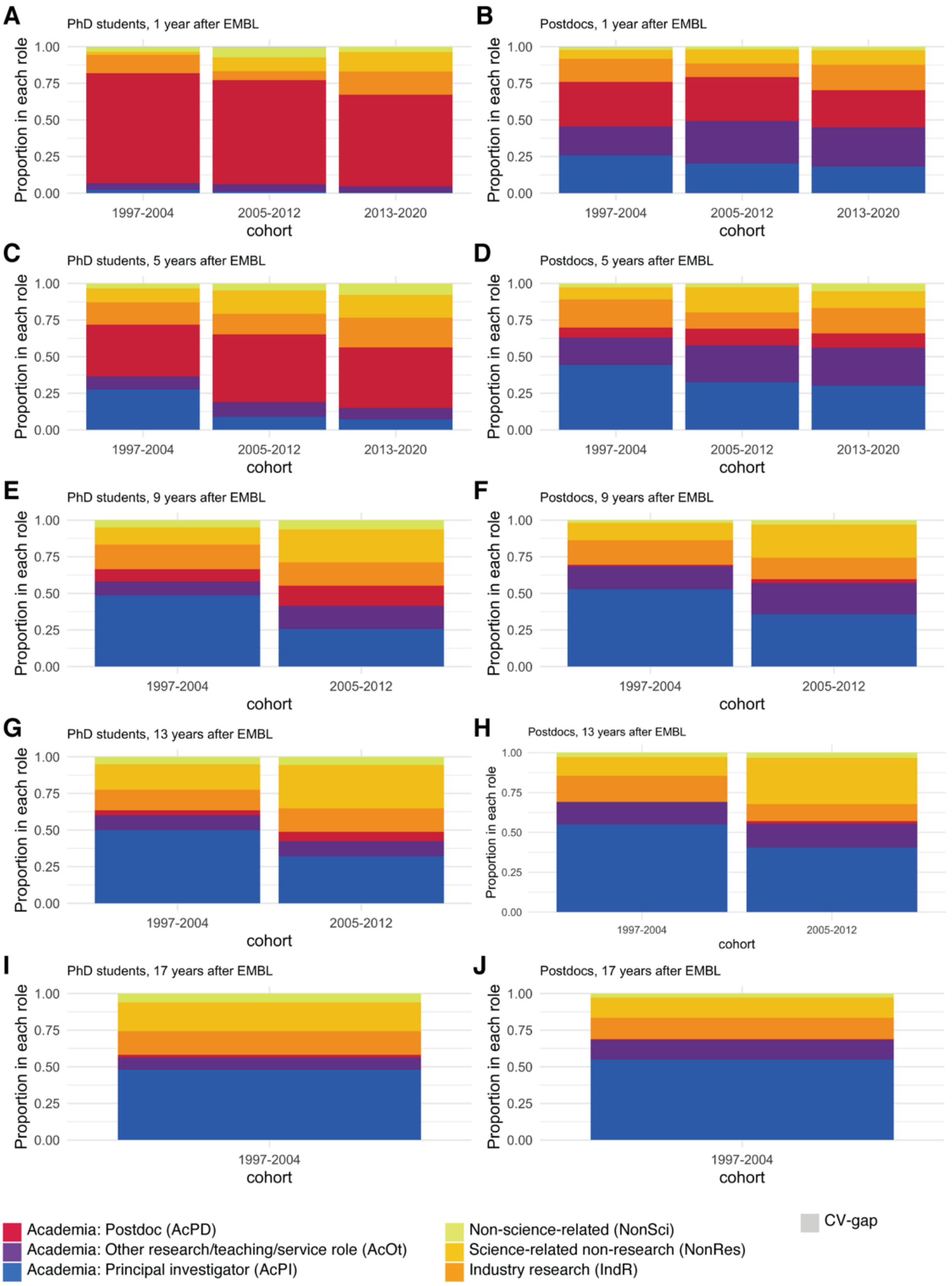

**Supplementary Figure 2: Role of alumni at specific timepoints after EMBL, by cohort and alumni type**

Column charts showing the percentage of alumni from each cohort in different types of role at specific timepoints after EMBL by cohort. Only alumni with a detailed career path available are included (n=1626). Fellows who have not yet reached the critical time point (e.g. for 5-years, those who left EMBL only four or less years ago) are omitted. **(A)** PhD alumni 1-year after EMBL. **(B)** Postdoc alumni 1-year after EMBL. **(C)** PhD alumni 5-years after EMBL. **(D)** Postdoc alumni 5-years after EMBL. **(E)** PhD alumni 9-years after EMBL. **(F)** Postdoc alumni 9-years after EMBL. **(G)** PhD alumni 13-years after EMBL **(H)** Postdoc alumni 13-years after EMBL. **(I)** PhD alumni 17-years after EMBL **(J)** Postdoc alumni 17-years after EMBL.

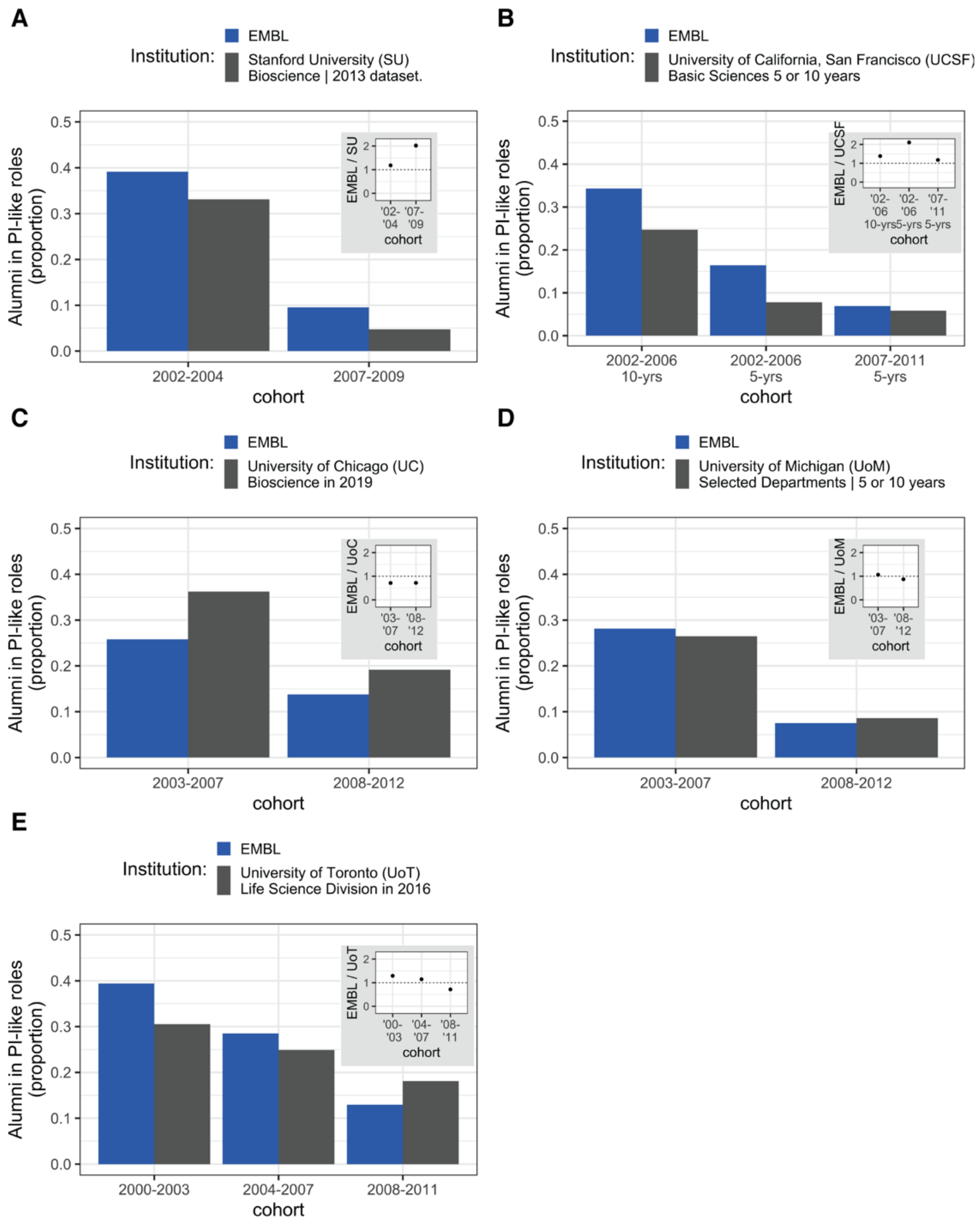

**Supplementary Figure 3: Proportion of alumni from other institutions found in tenure-track (TT) faculty-roles.**

Column charts showing cohort-based data on the proportion of PhD alumni from other institutions (grey) in tenure-track or tenure-stream positions compared to equivalent data on the proportion of PIs from this study for EMBL PhD alumni (blue columns). The inset shows the fold-difference between the EMBL proportion and proportion from the other institution for each cohort. Detailed information about the comparison groups can be found in Supplementary Table 4. Very recent cohorts have been omitted, as for these groups most PhD alumni are still in postdoc roles. The figures provided are the numbers known to be in a PI-like role at the relevant time-point as a proportion of all PhD students from the specified programme and PhD defence years. This includes students whose role was unknown at that time point. **(A)** Proportion of PhD alumni from 2002-2006, 2007-2011 or 2012-2016 in tenure-track (TT) faculty roles 5- or 10-years after PhD from University of California San Francisco (UCSF) (65) **(B)** Proportion of University of Chicago Bioscience Division PhD alumni from 2005-2009 or 2010-2014 in faculty roles in 2019 (66) **(C)** Proportion of alumni from programmes from the University of Michigan with overlap to EMBL research areas (see Supplementary Table 5) from 2003-2007 and 2008-2012 in tenure-track (TT) faculty roles 5 or 10 years after PhD. (67) **(D)** Proportion of alumni from programmes from the University of Toronto Life Sciences division from 2000-2003, 2004-2007 and 2008-2011 in tenure-stream faculty roles in 2016. (32, 68) **(E)** Column charts showing the proportion of EMBL PhD alumni who were found as principal investigators compared to a second set of cohort-based data from Stanford University (US) for tenure-track (TT) faculty collated in 2013 (64).

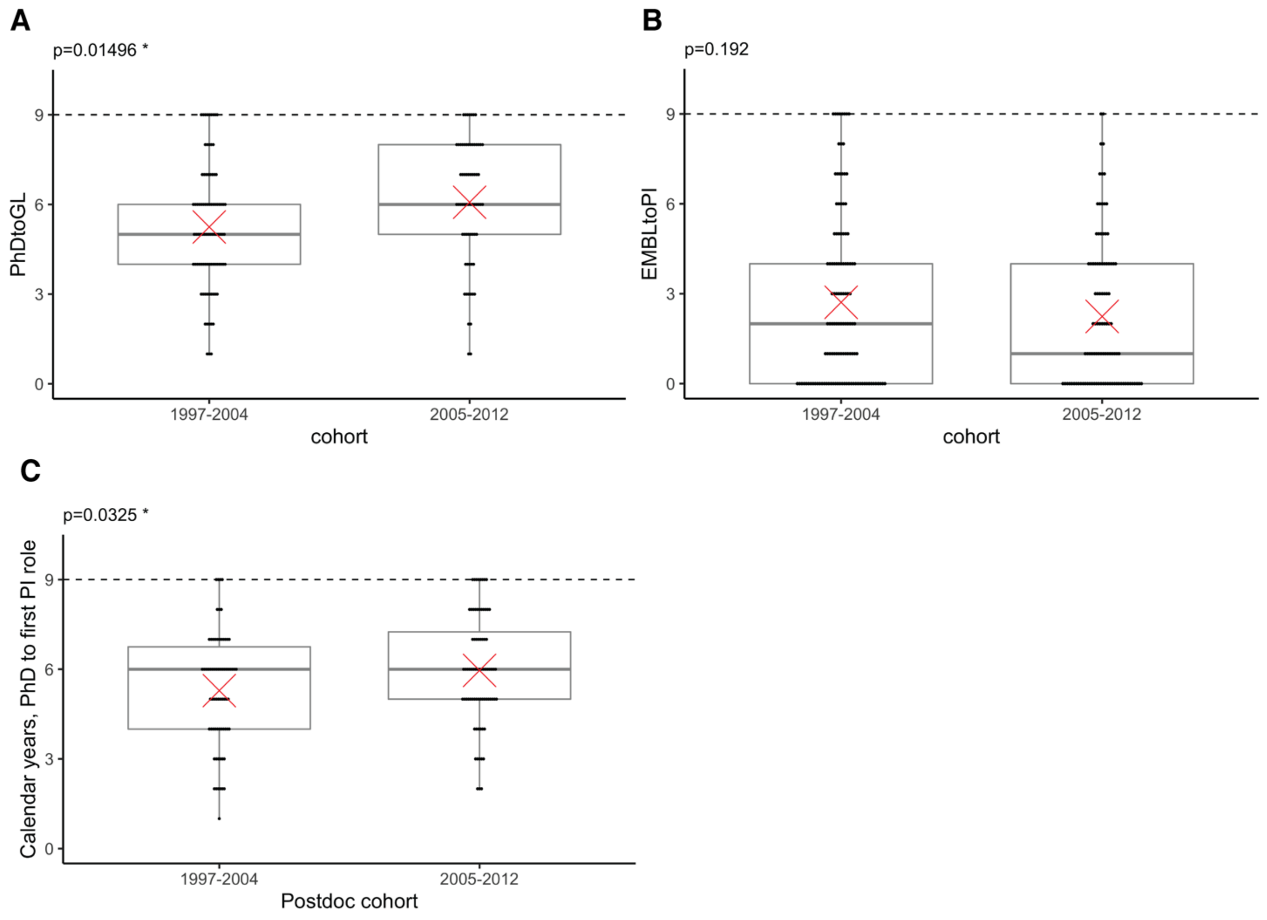

**Supplementary Figure 4: Comparison of time to first principal investigator role by cohort**
**(A)** Box-plot with overlaid dot-plot showing the distribution in calendar years between PhD conferral and start of first PI role for PhD alumni who defended in 1997-2012 who became a principal investigator 0 to 9 calendar years after their EMBL PhD defence, and for whom we have a detailed career path (n=157). The mean value is indicated as a red cross and the p-values calculated using Welch's t-test **(B)** Box-plot with overlaid dot-plot showing the distribution in calendar years between completion of an EMBL postdoc and start of first PI role for postdoc alumni who completed their postdoc in 1997-2012 who became a principal investigator 0 to 9 calendar years after their EMBL postdoc, and for whom we have a detailed career path (n=218). The mean value is indicated as a red cross and the p-values calculated using Welch's t-test. **(C)** As (B) but for postdoc alumni for whom we were able to identify a PhD conferral year (n=146).

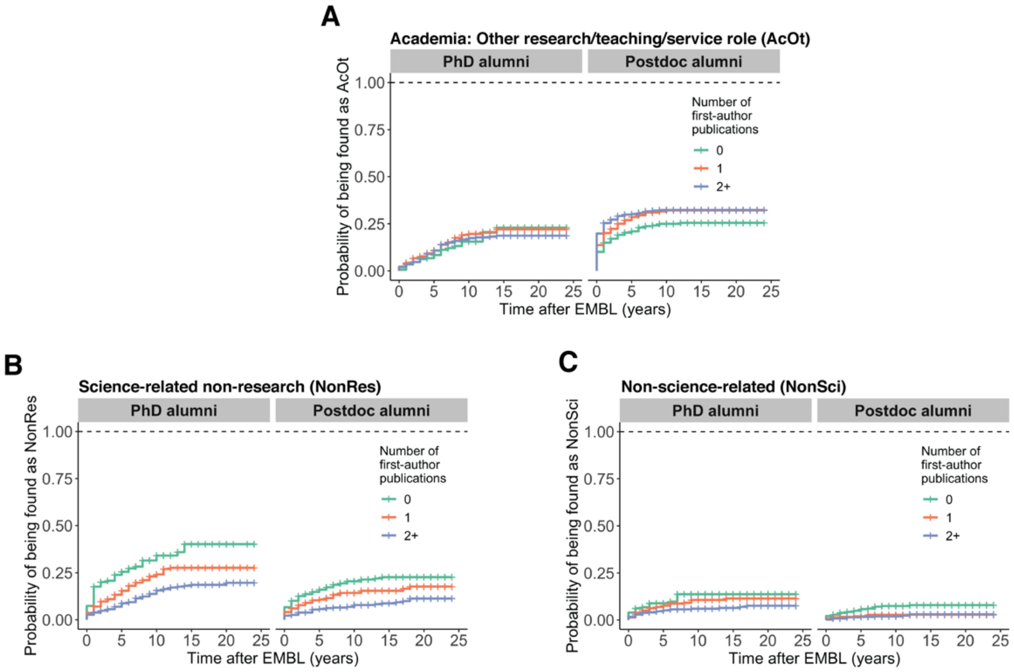

**Supplementary Figure 5: Entry into non-PI career areas by number of first author publications**
Kaplan Meier-curves for probability of entering specific non-PI career areas stratified by number of first-author publications from research completed at EMBL.

#### Supplementary Table 1: Descriptive statistics for the key data set used for figures

| Variable | Stats / Values | Freqs (% of Valid) | Valid | Missing | Description |
|---|---|---|---|---|---|
| unique_ID<br>[character] | [ 2284 unique values ] | | 2284<br>(100.0%) | 0<br>(0.0%) | unique ID number (F + unique random number) |
| type_pre_postdoc<br>[character] | 1. postdoc<br>2. predoc | 1315 (57.6%)<br>969 (42.4%) | 2284<br>(100.0%) | 0<br>(0.0%) | predoc (PhD student) or postdoc |
| gender<br>[character] | 1. f<br>2. m | 907 (39.7%)<br>1377 (60.3%) | 2284<br>(100.0%) | 0<br>(0.0%) | (f= female, m=male) |
| nationality<br>[character] | 1. [coding suppressed]<br>2. [coding suppressed]<br>3. [coding suppressed]<br>4. [coding suppressed]<br>5. [coding suppressed]<br>6. [coding suppressed]<br>7. [coding suppressed]<br>8. [coding suppressed]<br>9. [coding suppressed]<br>10. [coding suppressed]<br>[ 82 others ] | 486 (21.3%)<br>239 (10.5%)<br>206 ( 9.0%)<br>152 ( 6.7%)<br>121 ( 5.3%)<br>95 ( 4.2%)<br>66 ( 2.9%)<br>59 ( 2.6%)<br>58 ( 2.5%)<br>56 ( 2.5%)<br>746 (32.7%) | 2284<br>(100.0%) | 0<br>(0.0%) | Pseudomized Nationality of ECR |
| phd_year_if_known<br>[integer] | Mean (sd) : 2007.6 (7.1)<br>min < med < max:<br>1984 < 2008 < 2020<br>IQR (CV) : 11 (0) | 36 distinct values | 1870<br>(81.9%) | 414<br>(18.1%) | Year PhD awarded (if known) |
| from.year<br>[integer] | Mean (sd) : 2005.8 (6.8)<br>min < med < max:<br>1986 < 2007 < 2019<br>IQR (CV) : 12 (0) | 33 distinct values | 2284<br>(100.0%) | 0<br>(0.0%) | Start year as PhD student or postdoc |
| to_year<br>[integer] | Mean (sd) : 2009.4 (6.8)<br>min < med < max:<br>1997 < 2010 < 2020<br>IQR (CV) : 11 (0) | 24 distinct values | 2284<br>(100.0%) | 0<br>(0.0%) | Year of PhD defence or end of postdoc contract |
| cohort<br>[character] | 1. 1997-2004<br>2. 2005-2012<br>3. 2013-2020 | 625 (27.4%)<br>763 (33.4%)<br>896 (39.2%) | 2284<br>(100.0%) | 0<br>(0.0%) | Cohorts based on year_to (year of PhD defense or year they left the EMBL Postdoc Programme) |
| completeness<br>[character] | 1. Complete CV<br>2. Current only<br>3. No info<br>4. Partial CV | 1626 (71.2%)<br>274 (12.0%)<br>175 ( 7.7%)<br>209 ( 9.2%) | 2284<br>(100.0%) | 0<br>(0.0%) | Complete CV is for those who we could trace a detailed career path for, with max. two 1-year career breaks. |
| position_1<br>[character] | 1. AcFac<br>2. AcOt<br>3. AcPD<br>4. IndR<br>5. laterPI<br>6. NonRes<br>7. NonSci<br>8. unknown | 198 ( 8.7%)<br>280 (12.3%)<br>827 (36.2%)<br>217 ( 9.5%)<br>113 ( 4.9%)<br>146 ( 6.4%)<br>56 ( 2.5%)<br>447 (19.6%) | 2284<br>(100.0%) | 0<br>(0.0%) | Type of role they had at particular timepoints: last (position in 2021) and positions at 1,3,5 etc years after pre/postdoc contract ends)).<br>8 classifications were used:<br>· AcFac [principal investigator in academia]<br>· AcOt [other academic research / teaching role]<br>· AcPD [academic postdoc]<br>· IndR [Industry Research]<br>· NonRes [science related non-research professions]<br>· NonSci [non scientific professions]<br>· Later PI [position unknown at this timepoint, but became PI]<br>· unknown<br>The cell is empty if the ECR has not yet reached this timepoint |
| position_5<br>[character] | 1. (Empty string)<br>2. AcFac<br>3. AcOt<br>4. AcPD<br>5. IndR<br>6. laterPI<br>7. NonRes<br>8. NonSci<br>9. unknown | 431 (18.9%)<br>350 (15.3%)<br>239 (10.5%)<br>357 (15.6%)<br>223 ( 9.8%)<br>106 ( 4.6%)<br>187 ( 8.2%)<br>60 ( 2.6%)<br>331 (14.5%) | 2284<br>(100.0%) | 0<br>(0.0%) | |
| position_9<br>[character] | 1. (Empty string)<br>2. AcFac<br>3. AcOt<br>4. AcPD<br>5. IndR<br>6. laterPI<br>7. NonRes<br>8. NonSci<br>9. unknown | 896 (39.2%)<br>397 (17.4%)<br>180 ( 7.9%)<br>74 ( 3.2%)<br>165 ( 7.2%)<br>81 ( 3.5%)<br>183 ( 8.0%)<br>43 ( 1.9%)<br>265 (11.6%) | 2284<br>(100.0%) | 0<br>(0.0%) | |

| | | | | | |
|---|---|---|---|---|---|
| position_13 [character] | 1. (Empty string)<br>2. AcFac<br>3. AcOt<br>4. AcPD<br>5. IndR<br>6. laterPI<br>7. NonRes<br>8. NonSci<br>9. unknown | 1287 (56.3%)<br>332 (14.5%)<br>94 ( 4.1%)<br>19 ( 0.8%)<br>107 ( 4.7%)<br>69 ( 3.0%)<br>150 ( 6.6%)<br>31 ( 1.4%)<br>195 ( 8.5%) | 2284 (100.0%) | 0 (0.0%) | |
| position_17 [character] | 1. (Empty string)<br>2. AcFac<br>3. AcOt<br>4. AcPD<br>5. IndR<br>6. laterPI<br>7. NonRes<br>8. NonSci<br>9. unknown | 1659 (72.6%)<br>232 (10.2%)<br>57 ( 2.5%)<br>5 ( 0.2%)<br>68 ( 3.0%)<br>48 ( 2.1%)<br>78 ( 3.4%)<br>22 ( 1.0%)<br>115 ( 5.0%) | 2284 (100.0%) | 0 (0.0%) | |
| timepointLast [character] | 1. AcFac<br>2. AcOt<br>3. AcPD<br>4. IndR<br>5. NonRes<br>6. NonSci<br>7. unknown | 636 (27.8%)<br>383 (16.8%)<br>244 (10.7%)<br>332 (14.5%)<br>349 (15.3%)<br>91 ( 4.0%)<br>249 (10.9%) | 2284 (100.0%) | 0 (0.0%) | |
| was_GL [character] | 1. (Empty string)<br>2. y | 1599 (70.0%)<br>685 (30.0%) | 2284 (100.0%) | 0 (0.0%) | y, if ECR held at least one "AcFac" position |
| EMBLtoPI [integer] | Mean (sd) : 3.8 (3.5)<br>min < med < max:<br>-1 < 3 < 18<br>IQR (CV) : 5 (0.9) | 19 distinct values | 539 (23.6%) | 1745 (76.4%) | Number of calendar years between completing their PhD/postdoc at EMBL and the year they started their first principal investigator role (if known). |
| PhDtoGL [integer] | Mean (sd) : 6.8 (3.3)<br>min < med < max:<br>1 < 6 < 21<br>IQR (CV) : 3 (0.5) | 21 distinct values | 489 (21.4%) | 1795 (78.6%) | Number of calendar years between their PhD and the year they started their first principal investigator role (if known). |
| new_AcFac_previous [character] | 1. (Empty string)<br>2. AcOt<br>3. AcPD<br>4. AcPhD<br>5. IndR<br>6. NonRes<br>7. NonSci | 1745 (76.4%)<br>111 ( 4.9%)<br>406 (17.8%)<br>6 ( 0.3%)<br>14 ( 0.6%)<br>1 ( 0.0%)<br>1 ( 0.0%) | 2284 (100.0%) | 0 (0.0%) | For alumni who held an 'AcFac' position, this cell lists the classification of the position held before the individual's first 'AcFac' role. |
| CV_wasAcOt [character] | 1. (Empty string)<br>2. y | 1617 (70.8%)<br>667 (29.2%) | 2284 (100.0%) | 0 (0.0%) | y, if ECR held at least one "AcOt" position |
| CV_PhDtoAcOt [integer] | Mean (sd) : 5.6 (3.1)<br>min < med < max:<br>-1 < 6 < 17<br>IQR (CV) : 4 (0.6) | 17 distinct values | 396 (17.3%) | 1888 (82.7%) | Number of calendar years between their PhD completion and the year they started their first AcOt role (if known/applicable). |
| CV_EMBLtoAcOt [integer] | Mean (sd) : 2.4 (3.1)<br>min < med < max:<br>-1 < 1 < 14<br>IQR (CV) : 4 (1.3) | 16 distinct values | 477 (20.9%) | 1807 (79.1%) | Number of calendar years between completing their PhD/postdoc at EMBL and the year they started their year they started their first AcOt role (if known/applicable) |

| | | | | | |
|---|---|---|---|---|---|
| new_AcOt_previous [character] | 1. (Empty string)<br>2. AcFac<br>3. AcPD<br>4. AcPhD<br>5. IndR<br>6. NonRes<br>7. NonSci | 1807 (79.1%)<br>7 ( 0.3%)<br>393 (17.2%)<br>38 ( 1.7%)<br>21 ( 0.9%)<br>12 ( 0.5%)<br>6 ( 0.3%) | 2284 (100.0%) | 0 (0.0%) | For alumni who held an ‘AcOt’ position, this cell lists the classification of the position held before the individual’s first ‘AcOt’ role. |
| CV_wasIndR [character] | 1. (Empty string)<br>2. y | 1811 (79.3%)<br>473 (20.7%) | 2284 (100.0%) | 0 (0.0%) | y, if ECR held at least one “IndR” position |
| CV_PhDtoIndR [integer] | Mean (sd) : 5 (4.2)<br>min < med < max:<br>-2 < 4 < 19<br>IQR (CV) : 5 (0.8) | 22 distinct values | 363 (15.9%) | 1921 (84.1%) | Number of calendar years between their PhD completion and the year they started their first IndR role (if known/applicable). |
| CV_EMBLtoIndR [integer] | Mean (sd) : 2.8 (3.7)<br>min < med < max:<br>-2 < 1 < 20<br>IQR (CV) : 4 (1.3) | 21 distinct values | 414 (18.1%) | 1870 (81.9%) | Number of calendar years between completing their PhD/postdoc at EMBL and the year they started their year they started their first IndR role (if known/applicable |
| new_IndR_previous [character] | 1. (Empty string)<br>2. AcFac<br>3. AcOt<br>4. AcPD<br>5. AcPhD<br>6. NonRes<br>7. NonSci | 1869 (81.8%)<br>23 ( 1.0%)<br>55 ( 2.4%)<br>234 (10.2%)<br>84 ( 3.7%)<br>15 ( 0.7%)<br>4 ( 0.2%) | 2284 (100.0%) | 0 (0.0%) | For alumni who held an ‘IndR’ position, this cell lists the classification of the position held before the individual’s first ‘IndR’ role. |
| CV_wasNonR [character] | 1. (Empty string)<br>2. y | 1857 (81.3%)<br>427 (18.7%) | 2284 (100.0%) | 0 (0.0%) | y, if ECR held at least one “NonR” position |
| CV_PhDtoNonRes [integer] | Mean (sd) : 5.3 (4.1)<br>min < med < max:<br>-1 < 5 < 22<br>IQR (CV) : 6 (0.8) | 20 distinct values | 314 (13.7%) | 1970 (86.3%) | Number of calendar years between their PhD completion and the year they started their first NonRes role (if known/applicable). |
| CV_EMBLtoNonRes [integer] | Mean (sd) : 3.9 (4.2)<br>min < med < max:<br>-7 < 2 < 20<br>IQR (CV) : 5.2 (1.1) | 23 distinct values | 364 (15.9%) | 1920 (84.1%) | Number of calendar years between completing their PhD/postdoc at EMBL and the year they started their year they started their first NonRes role (if known/applicable |
| new_SciR_previous [character] | 1. (Empty string)<br>2. AcFac<br>3. AcOt<br>4. AcPD<br>5. AcPhD<br>6. IndR<br>7. NonSci | 1920 (84.1%)<br>8 ( 0.4%)<br>37 ( 1.6%)<br>150 ( 6.6%)<br>70 ( 3.1%)<br>61 ( 2.7%)<br>38 ( 1.7%) | 2284 (100.0%) | 0 (0.0%) | y, if ECR held at least one “NonSci” position |
| CV_wasNonSci [character] | 1. (Empty string)<br>2. y | 2122 (92.9%)<br>162 ( 7.1%) | 2284 (100.0%) | 0 (0.0%) | For alumni who held an ‘NonSci’ position, this cell lists the classification of the position held before the individual’s first ‘NonSci’ role. |
| CV_PhDtoNonSci [integer] | Mean (sd) : 4.2 (4)<br>min < med < max:<br>-1 < 3 < 17<br>IQR (CV) : 6 (1) | 17 distinct values | 117 (5.1%) | 2167 (94.9%) | Number of calendar years between their PhD completion and the year they started their first NonSci role (if known/applicable). |

| | | | | | |
|---|---|---|---|---|---|
| CV_EMBLtoNonSci [integer] | Mean (sd) : 3.2 (3.7) min < med < max: -1 < 2 < 17 IQR (CV) : 4 (1.1) | 17 distinct values | 131 (5.7%) | 2153 (94.3%) | Number of calendar years between completing their PhD/postdoc at EMBL and the year they started their year they started their first NonSci role (if known/applicable |
| new_NonSci_previous [character] | 1. (Empty string)<br>2. AcFac<br>3. AcOt<br>4. AcPD<br>5. AcPhD<br>6. IndR<br>7. NonRes | 2153 (94.3%)<br>2 ( 0.1%)<br>8 ( 0.4%)<br>44 ( 1.9%)<br>33 ( 1.4%)<br>13 ( 0.6%)<br>31 ( 1.4%) | 2284 (100.0%) | 0 (0.0%) | For alumni who held a Non-research science-related 'NonRes ' position, this cell lists the classification of the position held before the individual's first 'NonRes' role. |
| PUBS_ALL_TOTAL [integer] | Mean (sd) : 4.5 (4.8) min < med < max: 0 < 3 < 64 IQR (CV) : 4 (1.1) | 37 distinct values | 2284 (100.0%) | 0 (0.0%) | **Total number of publications linked to EMBL in Web of Science that include this person as an author, excluding retracted publications and corrections.** |
| PUBS_ALL_JIF_count [integer] | Mean (sd) : 4.5 (4.8) min < med < max: 0 < 3 < 64 IQR (CV) : 5 (1.1) | 37 distinct values | 2284 (100.0%) | 0 (0.0%) | For each ECR, the number of EMBL publications for which a journal impact factor (JIF) is available. |
| PUBS_ALL_JIF_mean [numeric] | Mean (sd) : 10 (7.4) min < med < max: 0 < 8.2 < 74.7 IQR (CV) : 8.1 (0.7) | 1758 distinct values | 2043 (89.4%) | 241 (10.6%) | For each ECR, the average JIF for all of their EMBL publications |
| PUBS_ALL_JIF_SUM [numeric] | Mean (sd) : 51.6 (62.6) min < med < max: 0.7 < 31.7 < 685.5 IQR (CV) : 49.7 (1.2) | 1749 distinct values | 2022 (88.5%) | 262 (11.5%) | For each ECR, the sum of the JIFs for all of their EMBL publications |
| pubs_ALL_JIF_MAX [numeric] | Mean (sd) : 19.8 (15.1) min < med < max: 0.7 < 12.1 < 74.7 IQR (CV) : 29.8 (0.8) | 222 distinct values | 2022 (88.5%) | 262 (11.5%) | For each ECR, the highest JIF of their EMBL publications |
| pubs_All_pecentile_mean [numeric] | Mean (sd) : 65.1 (21.8) min < med < max: 0 < 67.8 < 100 IQR (CV) : 29.8 (0.3) | 2016 distinct values | 2043 (89.4%) | 241 (10.6%) | For each ECR, the mean of the "percentile in subject area" f for all their EMBL publications |
| pubs_All_percentile_MAX [numeric] | Mean (sd) : 86.4 (20.5) min < med < max: 0 < 94.8 < 100 IQR (CV) : 15 (0.2) | 1542 distinct values | 2043 (89.4%) | 241 (10.6%) | For each ECR, the highest value of "percentile in subject area" for all their EMBL publications |
| PUBS_ALL_CNCI_mean [numeric] | Mean (sd) : 2.8 (5.4) min < med < max: 0 < 1.6 < 88.8 IQR (CV) : 2 (1.9) | 1982 distinct values | 2043 (89.4%) | 241 (10.6%) | For each ECR, the mean of the "category normalized citation impact" for their EMBL publications |
| PUBS_ALL_CNCI_max [numeric] | Mean (sd) : 9 (28.7) min < med < max: 0 < 3 < 441.2 IQR (CV) : 5.1 (3.2) | 1537 distinct values | 2043 (89.4%) | 241 (10.6%) | For each ECR, the highest value of "CNCI" for their EMBL publications |

| | | | | | |
|---|---|---|---|---|---|
| pubs_All_published.first [integer] | Mean (sd) : 2.4 (1.6) min < med < max: 0 < 2 < 14 IQR (CV) : 2 (0.7) | 14 distinct values | 2047 (89.6%) | 237 (10.4%) | Number of calendar years between EMBL start year and the publication date of their earliest publication |
| pubs_All_published.last [integer] | Mean (sd) : 5.4 (2.4) min < med < max: 0 < 5 < 18 IQR (CV) : 3 (0.4) | 19 distinct values | 2047 (89.6%) | 237 (10.4%) | For each ECR, the number of calendar years between EMBL start year and the publication date of their most publication |
| pubs_All_published_authors_mean [numeric] | Mean (sd) : 8.7 (9.5) min < med < max: 1 < 6.7 < 201 IQR (CV) : 4.2 (1.1) | 470 distinct values | 2047 (89.6%) | 237 (10.4%) | For each ECR, the mean number of authors on their research articles |
| pubs_RAonly_TOTAL [integer] | Mean (sd) : 3.6 (3.9) min < med < max: 0 < 3 < 53 IQR (CV) : 4 (1.1) | 31 distinct values | 2284 (100.0%) | 0 (0.0%) | **Total number of research articles linked to EMBL in Web of Science that include this person as an author, excluding retracted publications and corrections.** |
| pubs_RAonly_JIF_count [integer] | Mean (sd) : 3.6 (3.9) min < med < max: 0 < 3 < 53 IQR (CV) : 4 (1.1) | 31 distinct values | 2284 (100.0%) | 0 (0.0%) | Number of research articles for which a JIF is available. . |
| pubs_RAonly_JIF_mean [numeric] | Mean (sd) : 11 (8.1) min < med < max: 0 < 8.9 < 74.7 IQR (CV) : 8.9 (0.7) | 1617 distinct values | 1981 (86.7%) | 303 (13.3%) | For each ECR, the mean of the JIF for their EMBL research articles |
| pubs_RAonly_JIF_SUM [numeric] | Mean (sd) : 46.3 (55.7) min < med < max: 1 < 27.9 < 645.7 IQR (CV) : 46.1 (1.2) | 1610 distinct values | 1966 (86.1%) | 318 (13.9%) | For each ECR, the sum of the JIFs for all of their EMBL research articles |
| pubs_RA_JIF_max [numeric] | Mean (sd) : 19.4 (14.8) min < med < max: 1 < 12.1 < 74.7 IQR (CV) : 29.8 (0.8) | 192 distinct values | 1966 (86.1%) | 318 (13.9%) | For each ECR, the highest JIF for their EMBL research articles |
| pubs_RAonly_percentile_mean [numeric] | Mean (sd) : 73.3 (18.9) min < med < max: 0 < 76.7 < 100 IQR (CV) : 24.6 (0.3) | 1945 distinct values | 1981 (86.7%) | 303 (13.3%) | For each ECR, the mean of the “percentile in subject area” for their EMBL research articles |
| pubs_RA_pecentile_MAX [numeric] | Mean (sd) : 87.5 (17.8) min < med < max: 0 < 94.8 < 100 IQR (CV) : 14.6 (0.2) | 1471 distinct values | 1981 (86.7%) | 303 (13.3%) | For each ECR, the highest value of “percentile in subject area” for their EMBL research articles |
| pubs_RA_CNCI_mean [numeric] | Mean (sd) : 3.1 (5.8) min < med < max: 0 < 1.8 < 109.3 IQR (CV) : 2.2 (1.9) | 1939 distinct values | 1981 (86.7%) | 303 (13.3%) | For each ECR, the mean of the “category normalized citation impact” for their EMBL research articles |

| | | | | | |
|---|---|---|---|---|---|
| pubs_RAonly_CNCI_max [numeric] | Mean (sd) : 8.1 (24.1) min < med < max: 0 < 2.9 < 425.9 IQR (CV) : 4.6 (3) | 1482 distinct values | 1981 (86.7%) | 303 (13.3%) | For each ECR, the highest value of "CNCI" for their EMBL research articles |
| pubs_RAonly_publishedfirst [integer] | Mean (sd) : 2.7 (1.7) min < med < max: 0 < 2 < 14 IQR (CV) : 3 (0.6) | 14 distinct values | 1985 (86.9%) | 299 (13.1%) | Number of calendar years between EMBL start year and the publication date of their earliest for research article |
| pubs_RAonly_publishedlast [integer] | Mean (sd) : 5.5 (2.4) min < med < max: 0 < 5 < 18 IQR (CV) : 3 (0.4) | 19 distinct values | 1985 (86.9%) | 299 (13.1%) | For each ECR, the number of calendar years between EMBL start year and the publication date of their most recent research article |
| pubs_RAonly_Authors_average [numeric] | Mean (sd) : 9.2 (9.9) min < med < max: 1 < 7 < 201 IQR (CV) : 5 (1.1) | 404 distinct values | 1985 (86.9%) | 299 (13.1%) | For each ECR, the mean number of authors on their research articles |
| pubs_FIRST_ra_only_TOTAL [integer] | Mean (sd) : 1.6 (1.7) min < med < max: 0 < 1 < 19 IQR (CV) : 2 (1.1) | 15 distinct values | 2284 (100.0%) | 0 (0.0%) | **Total number research articles linked to EMBL in Web of Science that include this person as first author or other lead author (co-first or last), excluding retracted publications and corrections** |
| pubs_FIRST_ra_only_JIF_count [integer] | Mean (sd) : 1.6 (1.7) min < med < max: 0 < 1 < 19 IQR (CV) : 2 (1.1) | 15 distinct values | 2284 (100.0%) | 0 (0.0%) | Number of first author research articles for which a JIF is available. |
| pubs_FIRST_ra_only_JIF_mean [numeric] | Mean (sd) : 10.5 (9.4) min < med < max: 0 < 8 < 64.8 IQR (CV) : 7.6 (0.9) | 966 distinct values | 1658 (72.6%) | 626 (27.4%) | For each ECR, the mean of the JIF for their first-author EMBL research articles |
| pubs_FIRST_ra_only_JIFsum [numeric] | Mean (sd) : 21.7 (21.9) min < med < max: 1 < 13.1 < 165.3 IQR (CV) : 20.8 (1) | 973 distinct values | 1633 (71.5%) | 651 (28.5%) | For each ECR, the sum of the JIFs for all of their EMBL first-author research articles |
| pubs_FIRST_RA_JIF_max [numeric] | Mean (sd) : 14.6 (13.1) min < med < max: 1 < 9.9 < 64.8 IQR (CV) : 10.3 (0.9) | 205 distinct values | 1633 (71.5%) | 651 (28.5%) | For each ECR, the highest JIF for their first-author EMBL research articles |
| pubs_FIRST_RA_pecentile_mean [numeric] | Mean (sd) : 72.3 (21.8) min < med < max: 0 < 76.8 < 100 IQR (CV) : 30.9 (0.3) | 1597 distinct values | 1658 (72.6%) | 626 (27.4%) | For each ECR, the mean of the "percentile in subject area" for their first-author EMBL research articles |
| pubs_FIRST_RA_pecentile_MAX [numeric] | Mean (sd) : 81.1 (21.6) min < med < max: 0 < 89.9 < 100 IQR (CV) : 24.8 (0.3) | 1488 distinct values | 1658 (72.6%) | 626 (27.4%) | For each ECR, the highest value of "percentile in subject area" for their first-author EMBL research articles |

| | | | | | |
|---|---|---|---|---|---|
| pubs_FIRST_RA_CNCI_mean [numeric] | Mean (sd) : 2.6 (4.9)<br>min < med < max:<br>0 < 1.5 < 92.7<br>IQR (CV) : 2 (1.9) | 1578 distinct values | 1658 (72.6%) | 626 (27.4%) | For each ECR, the mean of the "category normalized citation impact" for their first-author EMBL research articles |
| pubs_FIRST_RA_CNCI_max [numeric] | Mean (sd) : 4 (8.9)<br>min < med < max:<br>0 < 2 < 184.2<br>IQR (CV) : 3.1 (2.2) | 1482 distinct values | 1658 (72.6%) | 626 (27.4%) | For each ECR, the highest value of "CNCI" for their first-author EMBL research articles |
| pubs_FIRST_ra_only_publishedfirst [integer] | Mean (sd) : 3.3 (1.9)<br>min < med < max:<br>0 < 3 < 17<br>IQR (CV) : 2 (0.6) | 14 distinct values | 1666 (72.9%) | 618 (27.1%) | Number of calendar years between EMBL start year and the publication date of their earliest for research article |
| pubs_FIRST_ra_only_publishedlast [integer] | Mean (sd) : 4.8 (2)<br>min < med < max:<br>0 < 5 < 17<br>IQR (CV) : 2 (0.4) | 18 distinct values | 1666 (72.9%) | 618 (27.1%) | For each ECR, the number of calendar years between EMBL start year and the publication date of their most recent research article |
| pubs_FIRST_ra_only_Authors_mean [numeric] | Mean (sd) : 7 (6.4)<br>min < med < max:<br>1 < 5.6 < 105.5<br>IQR (CV) : 4 (0.9) | 153 distinct values | 1666 (72.9%) | 618 (27.1%) | For each ECR, the mean number of authors on their research articles |
| groupleader_seniority [character] | 1. (Empty string)<br>2. junior<br>3. senior | 14 ( 0.6%)<br>1078 (47.2%)<br>1192 (52.2%) | 2284 (100.0%) | 0 (0.0%) | Junior (GL not a senior scientist and had spent <10 years at EMBL when ECR left) / senior (senior scientist or ≥10 at EMBL when ECR left) |
| Group_pubs_ALL_TOTALrecords [integer] | Mean (sd) : 99.3 (118.3)<br>min < med < max:<br>0 < 63 < 688<br>IQR (CV) : 82 (1.2) | 98 distinct values | 2268 (99.3%) | 16 (0.7%) | Total of **pubs_ALL_TOTAL** for all ECRs in this study with this group leader [note: publications with more than one co-author counted multiple times] |
| Group_pubs_ALL_TOTALmean [numeric] | Mean (sd) : 4.6 (2.3)<br>min < med < max:<br>0.5 < 3.9 < 13.3<br>IQR (CV) : 2.2 (0.5) | 141 distinct values | 2266 (99.2%) | 18 (0.8%) | Average number of WOS indexed publications from ECRs in this study with this person's PI .- for each group [Group_pubs_ALL_TOTALrecords / [number of ECRs from the research group who were included in this study] |
| Group_pubs_ALL_TOTAL_wJIF [integer] | Mean (sd) : 99.2 (118.3)<br>min < med < max:<br>1 < 63 < 688<br>IQR (CV) : 82 (1.2) | 98 distinct values | 2266 (99.2%) | 18 (0.8%) | Number of publication records of ECRs in this group for which a JIF is available |
| Group_pubs_ALL_JIFmean [numeric] | Mean (sd) : 10 (3.8)<br>min < med < max:<br>1.5 < 9.9 < 25.2<br>IQR (CV) : 5 (0.4) | 213 distinct values | 2266 (99.2%) | 18 (0.8%) | Average JIF of papers with an author from this group (Total of pubs_ALL_JIF_SUM for all ECRs with this group leader / **Group_pubs_ALL_TOTAL_wJIF**). Note: papers with multiple authors are included multiple times in the calculation |
| Group_pubs_ALL_JIFtotalmean [numeric] | Mean (sd) : 45.9 (32.6)<br>min < med < max:<br>1.1 < 38.7 < 199.1<br>IQR (CV) : 29.8 (0.7) | 213 distinct values | 2266 (99.2%) | 18 (0.8%) | Average cumulative JIF for all papers for each ECR in this study with this person's PI (includes those from this person). [Total of pubs_ALL_JIF_SUM for all ECRs with this group leader / [number of ECRs from the research group who were included in this study]] Note: papers with two co-first authors are included twice in the calculation |

| | | | | | |
|---|---|---|---|---|---|
| Group_pubs_FIRST_ra_only_TOTALrecords [integer] | Mean (sd) : 34.3 (37.2) min < med < max: 0 < 23 < 209 IQR (CV) : 32 (1.1) | 54 distinct values | 2266 (99.2%) | 18 (0.8%) | Total of **pubs_FIRST_ra_only_TOTAL** for all ECRs with this group leader [note: publications with multiple co-first authors counted multiple times] |
| Group_pubsFIRST_ra_only_TOTALmean [numeric] | Mean (sd) : 1.6 (0.7) min < med < max: 0 < 1.4 < 6 IQR (CV) : 0.9 (0.4) | 107 distinct values | 2266 (99.2%) | 18 (0.8%) | Average number of first author research articles from ECRs in this study with this person's PI (includes those from this person) [**Group_pubs_FIRST_ra_only_TOTALrecords** / [number of ECRs from the research group who were included in this study]] |
| Group_pubs_FIRST_ra_only_TOTALrecords_wJIF [integer] | Mean (sd) : 34.4 (37.4) min < med < max: 1 < 23 < 210 IQR (CV) : 32 (1.1) | 53 distinct values | 2259 (98.9%) | 25 (1.1%) | Total number of publication records for all ECRs with this group leader that have a JIF [note: publications with multiple co-firsts counted multiple times] |
| Group_pubsFIRST_RAonly_JIFmean [numeric] | Mean (sd) : 10.2 (4.2) min < med < max: 1.4 < 9.8 < 25.7 IQR (CV) : 6 (0.4) | 205 distinct values | 2259 (98.9%) | 25 (1.1%) | Average JIF of research artilces with a first author from this group (Total of pubs_FIRST_ra_only_JIF_SUM for all ECRs with this group leader, divided by **Group_pubs_FIRST_ra_only_TOTALrecords**). Note: papers with two co-first authors are included twice in the calculation |
| Group_pubsFIRST_ra_only_JIFtotalmean [numeric] | Mean (sd) : 15.6 (8.4) min < med < max: 0 < 15.1 < 49.8 IQR (CV) : 10.6 (0.5) | 208 distinct values | 2266 (99.2%) | 18 (0.8%) | Average cumulative JIF for first author research articles for each ECR in this study with this person's PI (includes those from this person). [Total of pubs_FIRST_ra_only_JIF_SUM for all ECRs with this group leader / [number of ECRs from the research group who were included in this study]] Note: papers with two co-first authors are included twice in the calculation |
| Group_pubs_FIRST_ra_only_UNIQUE_pubs [integer] | Mean (sd) : 31.9 (33.8) min < med < max: 1 < 22 < 189 IQR (CV) : 32 (1.1) | 50 distinct values | 2259 (98.9%) | 25 (1.1%) | Number of unique research articles with an ECR from our study in this group as a first author (papers with 2 ifrst authors from same group counted once) |
| Group_pubs_FIRST_ra_only_UNIQUE_pubs_wJIF [integer] | Mean (sd) : 31.8 (33.8) min < med < max: 1 < 22 < 189 IQR (CV) : 32 (1.1) | 48 distinct values | 2259 (98.9%) | 25 (1.1%) | Number of unique research articles with an ECR from our study in this group as a first author that have a JIF |
| Group_pubs_FIRST_ra_only_JIF_mean_alternative [numeric] | Mean (sd) : 10.1 (4.2) min < med < max: 1.4 < 9.5 < 28.5 IQR (CV) : 5.6 (0.4) | 205 distinct values | 2259 (98.9%) | 25 (1.1%) | Average JIF for the papers in Group_pubs_FIRST_ra_only_UNIQUE_pubs Note: papers with two co-first authors are included ONCE in the calculation |
| Group_pubs_FIRST_ra_only_AUTHORmean [numeric] | Mean (sd) : 6.7 (3.2) min < med < max: 2 < 6 < 47 IQR (CV) : 2.5 (0.5) | 153 distinct values | 2259 (98.9%) | 25 (1.1%) | Average number of authors (including those outside this study) on the papers in Group_pubs_FIRST_ra_only_UNIQUE_pubs |

| | | | | | |
|---|---|---|---|---|---|
| Group_pubs_FIRST_ra_only_CNCImean [numeric] | Mean (sd) : 2.5 (2.1)<br>min < med < max:<br>0 < 2.1 < 26.7<br>IQR (CV) : 1.7 (0.8) | 206 distinct values | 2259<br>(98.9%) | 25<br>(1.1%) | Average CNCI for papers on the papers in Group_pubs_FIRST_ra_only_UNIQUE_pubs Note: papers with two co-first authors from this group are included ONCE in the calculation |
| Group_pubs_FIRST_ra_only_PERCENTILEmean [numeric] | Mean (sd) : 71.7 (10.5)<br>min < med < max:<br>0 < 72.8 < 99.5<br>IQR (CV) : 13 (0.1) | 206 distinct values | 2259<br>(98.9%) | 25<br>(1.1%) | Average percentile in subject area on the papers in Group_pubs_FIRST_ra_only_UNIQUE_pubs Note: papers with two co-first authors are included ONCE in the calculation |
| Group_pubsFIRST_RAonly_publishedfirst_same_type [numeric] | Mean (sd) : 3.2 (1.3)<br>min < med < max:<br>0 < 3 < 9.5<br>IQR (CV) : 1.7 (0.4) | 86 distinct values | 2122<br>(92.9%) | 162<br>(7.1%) | Average of pubs_FIRST_ra_only_publishedfirst for ECRs of the same type from this group in our study (e.g. if type_pre_postdoc = predoc, averages PhD students in this group (postdocs publish faster)) |

#### Supplementary Table 2: current workforce sector (vertical) and career type (horizontal) for all alumni in this study with a known sector and type of position (n=2031)

Excluding four alumni with a known type of position but where the sector of the position was unclear, and 249 alumni whose position we could not identify, are excluded from this analysis.

| | Research / Teaching | Science related non-research | Non-science related | Grand Total |
|---|---|---|---|---|
| Academia | 62.2% | 3.3% | 0.0% | **65.5%** |
| Government | 0.0% | 1.3% | 0.2% | **1.6%** |
| Industry / for-profit | 16.2% | 10.2% | 3.9% | **30.3%** |
| Non-profit | 0.1% | 1.2% | 0.1% | **1.4%** |
| Other | 0.7% | 0.5% | 0.0% | **1.2%** |
| | **79.2%** | **16.5%** | **4.3%** | **100.0%** |

* note – the work-sectors primary research and primarily teaching from the UCOT Exp2 taxonomy have been merged

#### Supplementary Table 3: career type and current job functions (n=2031)

Excluding four alumni with a known type of position but where the sector of the position was unclear, and 249 alumni whose position we could not identify, are excluded from this analysis.

| *Career type and job function* | *% of found alumni* |
|---|---|
| **Research / Teaching** | **79.3%** |
| Independent research group leader[#] | 31.3% |
| Other academic position[^] | 18.9% |
| Postdoctoral[%] | 12.4% |
| R & D scientist[§] | 15.0% |
| Business dev., consulting, & strategic alliances[*] | 0.2% |
| Entrepreneurship[°] | 0.7% |
| Science education and outreach | 0.7% |
| **Science related non-research** | **16.4%** |
| Administration and training | 3.4% |
| Business dev., consulting, & strategic alliances | 2.9% |
| Clinical research management | 0.6% |
| Clinical services / public health | 0.5% |
| Data science, analytics, software engineering[$] | 1.4% |
| Entrepreneurship | 0.2% |
| Healthcare provider | 0.4% |
| Intellectual property and law | 1.3% |
| Other | 0.3% |
| Regulatory affairs | 0.6% |
| Sales and marketing | 0.5% |
| Science education and outreach | 0.3% |
| Science policy and government affairs | 0.1% |
| Science writing and communication | 1.8% |
| Technical support and product development | 2.2% |
| **Non-science related** | **4.3%** |
| Administration and training | 0.1% |
| Business dev., consulting, & strategic alliances | 1.0% |
| Data science, analytics, software engineering[$] | 2.1% |
| Entrepreneurship | 0.3% |
| Other | 0.7% |

### includes those leading an academic research team with financial and scientific independence – evidenced by job title as 'group leader', 'professor', 'associate professor' or 'tenure-track assistant professor'. Where the status was unclear from the job title, we classified as independent research group leaders if one of the following criteria was fulfilled: a) they appear to directly supervise students/postdocs (based on hierarchy shown on website), b) they have published a last author publication from the current position c) their group website or CV indicates that they have a grant (not just a personal merit fellowship) as a principle investigator.
* includes director-level senior management roles overseeing the scientific direction & research of a company with R&D focus, e.g. CSOs in biotech start-ups
% 244 in academic sector and 8 in for-profit companies
° founders of companies whose primary focus is R&D (including contract research)
^ this function differs from the published schema; it includes academic research, scientific services or teaching staff e.g. research staff, teaching faculty and staff, technical directors, research infrastructure engineers
§ this function differs from the published schema; it includes alumni carrying out or overseeing scientific research in industry as group leaders, research staff, technical directors and non-directorship research leadership roles, including alumni who appear to be working in computational biology roles of a pharma, biotech, contract research or similar company regardless of job title i.e. including data science roles that appear to be related to analysis of research-related data.
[$] not including computational biology roles linked to R&D functions.

#### Supplementary table 4: Comparison datasets included in the comparisons in Figure 2B and Supplementary Figure 4

| Figure | Institution | Field(s) | Source | PI-like role defined as | Cohorts | Timing | Percentage of alumni in PI-like roles | Cohort size | Equivalent EMBL data | EMBL cohort size |
|---|---|---|---|---|---|---|---|---|---|---|
| S5A | University of California, San Francisco | Basic health science | https://graduate.ucsf.edu/program-statistics<br>Last accessed 9 Sept 2021 | Tenure-track faculty | 2002-2006 | 10 ys out<br>5 yrs out<br>1 yr out | 24.7%<br>7.8%<br>1.4% | 295 | 34.3%<br>16.4%<br>1.5% | 201 |
| | | | | | 2007-2011 | 5 yrs out<br>1 yr out | 5.9%<br>1.6% | 512 | 6.9%<br>1.8% | 217 |
| | | | | | 2012-2016 | 1 yr out | 1.2% | 521 | 0.4% | 246 |
| S5 B | University of Chicago | Bioscience | https://biosciences.uchicago.edu/after-uchicago/outcomes<br>Last accessed 20 Aug 2021 | Faculty incl. nonTT | 2005-2009<br>2010-2014<br>2015-2019 | in 2019 | 36.2%<br>19.2%<br>5.3% | 338<br>339<br>312 | 25.8%<br>13.8%<br>2.1% | 217<br>225<br>234 |
| S5 C | University of Michigan | Life sciences[1] | https://secure.rackham.umich.edu/academic_information/program_statistics/doctoral/<br>Data downloaded / compiled 9 Nov 2018 | Tenure-track faculty | 2003-2007<br>2008-2012<br>2013-2017 | 10 ys out<br>5 yrs out<br>1 yr out | 26.5%<br>8.6%<br>1.3% | 170<br>244<br>294 | 28.2%<br>7.5%<br>0.4% | 206<br>213<br>244 |
| S5D | Stanford University | Bioscience | https://irds.stanford.edu/data-findings/phd-jobs<br>Last accessed 17 Jan 2020 | Tenure-track | 2002-2004<br>2007-2009 | in 2013 | 33.1%<br>4.7% | 139<br>169 | 39.2%<br>9.6% | 120<br>136 |
| 2B | Stanford University | Bioscience | https://biosciences.stanford.edu/prospective-students/alumni-career-outcomes-dashboard/alumni-career-outcomes-by-cohort/<br>Last checked 10 Sept 21 | Faculty – research focus | 2000-2005<br>2006-2010<br>2011-2015<br>2016-2019 | in 2018[2] | 34.0%<br>26.8%<br>12.5%<br>2.8% | 426<br>471<br>503<br>472 | 37.1%<br>22.4%<br>10.7%<br>1.6% | 210<br>223<br>234<br>185 |
| S5E | University of Toronto | Life sciences division | Reithmeier, R., et al. (2019). " PLoS One 14(1): e0209898.<br>https://www.sgs.utoronto.ca/about/explore-our-data/10000-phds-project/<br>Data complied 10 Sep 2021 | Tenure stream | 2000-2003<br>2004-2007<br>2008-2011<br>2012-2015 | in 2016 | 30.5%<br>24.9%<br>18.1%<br>7.9% | 696<br>816<br>1028<br>1234 | 39.4%<br>28.5%<br>12.9%<br>1.6% | 132<br>172<br>170<br>193 |

Please note:

[1] The statistics from the following programmes, which overlap with EMBL's research areas, were combined: Bioinformatics, Cellular and Molecular Biology, Microbiology and Immunology, Molecular, Cellular and Developmental Biology, Cancer Biology, Biophysics, Biological Chemistry, Cell and Developmental Biology

[2] Except for the 2019 leavers: for this leaving year, the career destination in 2020 (position one calendar year after defence) was used.

- The most recent PhD cohorts and data on PhD destinations 1-year after graduation were not included in the plots in Figure 2B / S5. These values are shown in grey. We did not include this data as very few PhD alumni became a PI in the first years after leaving EMBL, consistent with almost all PhDs competing at least one postdoc position before gaining a PI role. A single alumnus/alumna more or less who become PIs at a very early stage will lead to large differences in the relative ratios that does not necessarily reflect a trend to fewer or more PIs in the long-term.
- One published study (Mathur, Cano (31)) was not included in these comparisons, as a high percentage of alumni (14% of all alumni) were in primarily teaching focused roles – many of these are likely to be included in the faculty numbers. Teaching-focused faculty roles may have different career dynamics to the careers of in research-focussed PI roles. The biomedical focus of the Mathur et al. study may also limit comparability compared to the basic life science focus of EMBL.

**Supplementary table 5: Hazard ratios of later cohorts compared with earlier reference cohorts, calculated by Cox regression model.**

A hazard ratio lower than 1 indicates decreased likelihood of entering that role.

| *Role of interest* | *Role at EMBL* | *Cohort* | *Reference cohort* | *Hazard ratio* | *95% Confidence Interval* | *p-value* | |
|---|---|---|---|---|---|---|---|
| ***PI*** | PhD students | **2005-2012** | **1997-2004** | **0.51** | **0.38 - 0.69** | **1.10E-05** | *** |
| | | **2013-2020** | **1997-2004** | **0.23** | **0.14 - 0.38** | **1.90E-08** | *** |
| | | 2013-2020 | 2005-2012 | 0.59 | 0.34 - 1.01 | 0.052 | |
| | Postdocs | **2005-2012** | **1997-2004** | **0.58** | **0.45 - 0.76** | **5.40E-05** | *** |
| | | **2013-2020** | **1997-2004** | **0.69** | **0.53 - 0.89** | **0.0054** | ** |
| | | 2013-2020 | 2005-2012 | 1.05 | 0.80 - 1.38 | 0.73 | |
| ***AcOt*** | PhD students | 2005-2012 | 1997-2004 | 1.14 | 0.78 - 1.68 | 0.49 | |
| | | 2013-2020 | 1997-2004 | 1.06 | 0.66 - 1.68 | 0.82 | |
| | | 2013-2020 | 2005-2012 | 1.03 | 0.67 - 1.59 | 0.89 | |
| | Postdocs | **2005-2012** | **1997-2004** | **1.72** | **1.16 - 2.10** | **0.0034** | ** |
| | | **2013-2020** | **1997-2004** | **1.09** | **1.29 - 2.31** | **0.00027** | *** |
| | | 2013-2020 | 2005-2012 | 1.72 | 0.85 - 1.40 | 0.52 | |
| ***IndR*** | PhD students | 2005-2012 | 1997-2004 | 1.11 | 0.76 - 1.60 | 0.6 | |
| | | **2013-2020** | **1997-2004** | **1.60** | **1.08 - 2.36** | **0.018** | * |
| | | **2013-2020** | **2005-2012** | **1.63** | **1.15 - 2.31** | **0.0064** | ** |
| | Postdocs | 2005-2012 | 1997-2004 | 0.72 | 0.49 - 1.04 | 0.08 | |
| | | **2013-2020** | **1997-2004** | **1.51** | **1.09 - 2.10** | **0.013** | * |
| | | **2013-2020** | **2005-2012** | **2.39** | **1.68 - 3.40** | **1.4E-06** | *** |
| ***NonRes*** | PhD students | **2005-2012** | **1997-2004** | **1.87** | **1.29 - 2.71** | **0.00089** | *** |
| | | **2013-2020** | **1997-2004** | **2.33** | **1.47 - 3.71** | **0.00036** | *** |
| | | 2013-2020 | 2005-2012 | 1.10 | 0.77 - 1.56 | 0.61 | |
| | Postdocs | **2005-2012** | **1997-2004** | **1.78** | **1.19 - 2.68** | **0.0055** | ** |
| | | **2013-2020** | **1997-2004** | **1.93** | **1.22 - 3.06** | **0.0053** | ** |
| | | 2013-2020 | 2005-2012 | 1.05 | 0.73 - 1.51 | 0.79 | |
| ***NonSci*** | PhD students | **2005-2012** | **1997-2004** | **2.02** | **1.10 - 3.73** | **0.024** | * |
| | | **2013-2020** | **1997-2004** | **2.21** | **1.12 - 4.38** | **0.022** | * |
| | | 2013-2020 | 2005-2012 | 1.06 | 0.63 - 1.76 | 0.84 | |
| | Postdocs | 2005-2012 | 1997-2004 | 1.44 | 0.67 - 3.08 | 0.35 | |
| | | **2013-2020** | **1997-2004** | **2.58** | **1.13 - 5.88** | **0.024** | * |
| | | 2013-2020 | 2005-2012 | 1.63 | 0.84 - 3.18 | 0.15 | |

#### Supplementary table 6: differences in publication statistics by PI status.

Values that are statistically significant (p<0.05) are highlighted with bold, and those mentioned in paper text are underlined.

| Publication statistics | Became PI? no | yes | sig~ | | Field name |
|---|---|---|---|---|---|
| *n* | 1599 | 685 | | | |
| For: ***all publications*** *from EMBL work* | | | | | |
| ***Number of publications*** | **3.9** | **6.1** | **<2e-16** | ******* | **PUBS_ALL_TOTAL** |
| ***Number of publications with a JIF*** | **3.9** | **6.1** | **<2e-16** | ******* | **PUBS_ALL_JIF_count** |
| ***Mean JIF*** | **9.6** | **10.9** | **1.2E-04** | ******* | **PUBS_ALL_JIF_mean** |
| ***Sum of JIFs*** | **42.1** | **71.9** | **<2e-16** | ******* | **PUBS_ALL_JIF_SUM** |
| ***Highest JIF*** | **17.8** | **24.0** | **<2e-16** | ******* | **pubs_ALL_JIF_MAX** |
| ***Mean of "percentile in subject area"*** | **63.1** | **69.6** | **4.8E-12** | ******* | **pubs_All_pecentile_mean** |
| ***Highest value of "percentile in subject area"*** | **83.9** | **91.7** | **<2e-16** | ******* | **pubs_All_percentile_MAX** |
| ***Mean CNCI*** | **2.6** | **3.1** | **4.6E-02** | ***** | **PUBS_ALL_CNCI_mean** |
| ***Highest value of "CNCI"*** | **8.1** | **11.0** | **2.8E-02** | ***** | **PUBS_ALL_CNCI_max** |
| ***Calendar years: EMBL start year -> earliest publ.*** | **2.5** | **2.1** | **4.5E-11** | ******* | **pubs_All_published.first** |
| *Calendar years: EMBL start year -> last* | 5.4 | 5.5 | 2.0E-01 | | pubs_All_published.last |
| ***Mean number of authors*** | **9.1** | **7.8** | **1.9E-03** | ****** | **pubs_All_published_authors_mean** |
| For: all research articles from EMBL work | | | | | |
| ***Number of research articles*** | **3.1** | **4.9** | **<2e-16** | ******* | **pubs_RAonly_TOTAL** |
| ***Number of research articles with a JIF*** | **3.1** | **4.9** | **<2e-16** | ******* | **pubs_RAonly_JIF_count** |
| ***Mean JIF*** | **10.5** | **11.9** | **3.3E-04** | ******* | **pubs_RAonly_JIF_mean** |
| ***Sum of JIFs*** | **38.4** | **62.8** | **8.8E-15** | ******* | **pubs_RAonly_JIF_SUM** |
| ***Highest JIF*** | **17.6** | **23.2** | **2.9E-14** | ******* | **pubs_RA_JIF_max** |
| ***Mean of "percentile in subject area"*** | **71.3** | **77.4** | **6.2E-14** | ******* | **pubs_RAonly_percentile_mean** |
| ***Highest value of "percentile in subject area"*** | **85.2** | **92.2** | **<2e-16** | ******* | **pubs_RA_pecentile_MAX** |
| ***Mean CNCI*** | **2.9** | **3.5** | **1.1E-02** | ***** | **pubs_RA_CNCI_mean** |
| ***Highest value of "CNCI"*** | **6.9** | **10.7** | **2.1E-03** | ****** | **pubs_RAonly_CNCI_max** |
| ***Calendar years: EMBL start-> earliest res.article*** | **2.8** | **2.3** | **7.6E-12** | ******* | **pubs_RAonly_publishedfirst** |
| *Calendar years: EMBL start year -> last* | 5.4 | 5.5 | 3.6E-01 | | pubs_RAonly_publishedlast |
| ***Mean number of authors*** | **9.7** | **8.2** | **6.5E-04** | ******* | **pubs_RAonly_Authors_average** |
| For: first-author research articles from EMBL work | | | | | |
| ***Number of 1st author research articles*** | **1.2** | **2.4** | **<2e-16** | ******* | **pubs_FIRST_ra_only_TOTAL** |
| ***Number of 1st author research articles with a JIF*** | **1.2** | **2.4** | **<2e-16** | ******* | **pubs_FIRST_ra_only_JIF_count** |
| ***Mean JIF*** | **9.7** | **12.1** | **7.9E-07** | ******* | **pubs_FIRST_ra_only_JIF_mean** |
| ***Sum of JIFs*** | **16.7** | **30.3** | **<2e-16** | ******* | **pubs_FIRST_ra_only_JIFsum** |
| ***Highest JIF*** | **12.3** | **18.5** | **<2e-16** | ******* | **pubs_FIRST_RA_JIF_max** |
| ***Mean of "percentile in subject area"*** | **69.3** | **77.6** | **7.0E-16** | ******* | **pubs_FIRST_RA_pecentile_mean** |
| ***Highest value of "percentile in subject area"*** | **77.1** | **88.1** | **<2e-16** | ******* | **pubs_FIRST_RA_pecentile_MAX** |
| ***Mean of CNCI*** | **2.2** | **3.3** | **2.6E-04** | ******* | **pubs_FIRST_RA_CNCI_mean** |
| ***Highest value of "CNCI"*** | **3.1** | **5.7** | **2.8E-06** | ******* | **pubs_FIRST_RA_CNCI_max** |
| ***Calendar years: EMBL start -> earliest 1st author*** | **3.6** | **2.8** | **7.5E-15** | ******* | **pubs_FIRST_ra_only_publishedfirst** |
| *Calendar years: EMBL start year -> last* | 4.8 | 4.8 | 9.3E-01 | | pubs_FIRST_ra_only_publishedlast |
| ***Mean number of authors*** | **7.3** | **6.4** | **6.0E-03** | ****** | **pubs_FIRST_ra_only_Authors_mean** |
| ***Number of co-author research articles*** | **1.9** | **2.5** | **1.5E-05** | ******* | **coauthor_research** |
| ***Number of non-research-articles*** | **0.8** | **1.2** | **2.3E-07** | ******* | **nonresearhc** |

~ statistical significance based on unpaired 2-tailed t-test *<0.05, **<0.01,***<0.001

**Supplementary table 7: Hazard ratios of those publishing multiple first author papers compared to those publishing 0 or 1 first author papers calculated by Cox regression model with time from EMBL.**

A hazard ratio lower than 1 indicates decreased likelihood of entering the specified role.

| *Entry into* | *Role at EMBL* | *Number of 1st-author* | *Reference # 1st-author* | *Hazard ratio* | *95% Confidence Interval* | *p-value* | |
|---|---|---|---|---|---|---|---|
| *PI* | PhD | 1 | 0 | 1.73 | 0.92 - 3.25 | 0.092 | |
| | | **2+** | **0** | **4.60** | **2.55 - 8.29** | **4E-07** | *** |
| | | **2+** | **1** | **2.66** | **1.91 - 3.71** | **7.1E-09** | *** |
| | postdoc | **1** | **0** | **3.24** | **2.23 - 4.72** | **8.1E-10** | *** |
| | | **2+** | **0** | **6.61** | **4.70 - 9.31** | **2.7E-27** | *** |
| | | **2+** | **1** | **2.04** | **1.59 - 2.60** | **1.4E-08** | *** |
| *AcOt* | PhD | 1 | 0 | 1.12 | 0.69 - 1.80 | 0.65 | |
| | | 2+ | 0 | 0.97 | 0.60 - 1.54 | 0.89 | |
| | | 2+ | 1 | 0.87 | 0.61 - 1.24 | 0.45 | |
| | postdoc | **1** | **0** | **1.34** | **1.01 - 1.79** | **0.044** | * |
| | | **2+** | **0** | **1.45** | **1.11 - 1.88** | **0.0062** | ** |
| | | 2+ | 1 | 1.09 | 0.84 - 1.41 | 0.5 | |
| *IndR* | PhD | 1 | 0 | 0.91 | 0.61 - 1.36 | 0.65 | |
| | | 2+ | 0 | 0.90 | 0.61 - 1.33 | 0.6 | |
| | | 2+ | 1 | 0.95 | 0.70 - 1.30 | 0.75 | |
| | postdoc | **1** | **0** | **0.72** | **0.51 - 1.00** | **0.048** | * |
| | | **2+** | **0** | **0.65** | **0.48 - 0.88** | **0.0062** | ** |
| | | 2+ | 1 | 0.90 | 0.64 - 1.28 | 0.57 | |
| *NonRes* | PhD | **1** | **0** | **0.61** | **0.43 - 0.86** | **0.0052** | ** |
| | | **2+** | **0** | **0.37** | **0.25 - 0.53** | **8E-08** | *** |
| | | **2+** | **1** | **0.60** | **0.43 - 0.83** | **0.0024** | ** |
| | postdoc | **1** | **0** | **0.69** | **0.48 - 0.98** | **0.038** | * |
| | | **2+** | **0** | **0.37** | **0.26 - 0.55** | **3.5E-07** | *** |
| | | **2+** | **1** | **0.54** | **0.36 - 0.82** | **0.0039** | ** |
| *NonSci* | PhD | 1 | 0 | 0.76 | 0.44 - 1.32 | 0.33 | |
| | | **2+** | **0** | **0.48** | **0.27 - 0.85** | **0.012** | * |
| | | 2+ | 1 | 0.61 | 0.37 - 1.02 | 0.06 | |
| | postdoc | **1** | **0** | **0.36** | **0.17 - 0.77** | **0.0081** | ** |
| | | **2+** | **0** | **0.33** | **0.17 - 0.65** | **0.0014** | ** |
| | | 2+ | 1 | 0.92 | 0.39 - 2.18 | 0.85 | |

**Supplementary table 8: differences in publication statistics by 'academic research / service / teaching staff' status.**

Values that are statistically significant (p<0.05) are highlighted with bold, and those mentioned in paper text are underlined.

| Publication statistics | Became AcOt? no | yes | sig~ | Field name |
|---|---|---|---|---|
| *Number of alumni* | 1617 | 667 | | |
| *For: **all publications** from EMBL work* | | | | |
| ***Number of publications*** | **4.4** | **5.0** | **6.9E-03 **** | **PUBS_ALL_TOTAL** |
| ***Number of publications with a JIF*** | **4.4** | **5.0** | **7.6E-03 **** | **PUBS_ALL_JIF_count** |
| ***Mean JIF*** | **10.5** | **9.0** | **2.7E-06 ***** | **PUBS_ALL_JIF_mean** |
| *Sum of JIFs* | 52.4 | 49.5 | 3.1E-01 | PUBS_ALL_JIF_SUM |
| ***Highest JIF*** | **20.4** | **18.4** | **8.0E-03 **** | **pubs_ALL_JIF_MAX** |
| ***Mean of "percentile in subject area"*** | **66.0** | **63.0** | **4.3E-03 **** | **pubs_All_pecentile_mean** |
| *Highest value of "percentile in subject area"* | 86.6 | 86.0 | 5.8E-01 | pubs_All_percentile_MAX |
| *Mean CNCI* | 2.9 | 2.6 | 3.9E-01 | PUBS_ALL_CNCI_mean |
| *Highest value of "CNCI"* | 9.1 | 8.8 | 8.3E-01 | PUBS_ALL_CNCI_max |
| *Calendar years: EMBL start year -> earliest* | 2.4 | 2.3 | 3.7E-01 | pubs_All_published.first |
| *Calendar years: EMBL start -> final publication* | 5.4 | 5.5 | 4.8E-01 | pubs_All_published.last |
| *Mean number of authors* | 8.7 | 8.8 | 8.7E-01 | pubs_All_published_authors_mean |
| *For: **all research articles** from EMBL work* | | | | |
| ***Number of research articles*** | **3.5** | **4.0** | **1.4E-02 *** | **pubs_RAonly_TOTAL** |
| ***Number of research articles with a JIF*** | **3.5** | **4.0** | **1.4E-02 *** | **pubs_RAonly_JIF_count** |
| ***Mean JIF*** | **11.4** | **9.9** | **3.7E-05 ***** | **pubs_RAonly_JIF_mean** |
| *Sum of JIFs* | 47.1 | 44.5 | 3.1E-01 | pubs_RAonly_JIF_SUM |
| ***Highest JIF*** | **20.0** | **18.1** | **6.4E-03 **** | **pubs_RA_JIF_max** |
| ***Mean of "percentile in subject area"*** | **73.9** | **72.0** | **4.4E-02 *** | **pubs_RAonly_percentile_mean** |
| *Highest value of "percentile in subject area"* | 87.6 | 87.3 | 7.4E-01 | pubs_RA_pecentile_MAX |
| *Mean CNCI* | 3.1 | 2.9 | 4.1E-01 | pubs_RA_CNCI_mean |
| *Highest value of "CNCI"* | 8.2 | 7.8 | 7.3E-01 | pubs_RAonly_CNCI_max |
| *Calendar years: EMBL start-> earliest res.article* | 2.7 | 2.6 | 1.9E-01 | pubs_RAonly_publishedfirst |
| *Calendar years: EMBL start -> final res.article* | 5.4 | 5.4 | 8.6E-01 | pubs_RAonly_publishedlast |
| *Mean number of authors* | 9.1 | 9.3 | 7.5E-01 | pubs_RAonly_Authors_average |
| *For: **first-author research articles** from EMBL work* | | | | |
| *Number of 1st author research articles* | 1.6 | 1.6 | 7.7E-01 | pubs_FIRST_ra_only_TOTAL |
| *No. of 1st author research articles w/ JIF* | 1.6 | 1.6 | 7.7E-01 | pubs_FIRST_ra_only_JIF_count |
| ***Mean JIF*** | **11.2** | **9.2** | **2.7E-05 ***** | **pubs_FIRST_ra_only_JIF_mean** |
| ***Sum of JIFs*** | **22.8** | **19.0** | **5.0E-04 ***** | **pubs_FIRST_ra_only_JIFsum** |
| ***Highest JIF*** | **15.4** | **12.7** | **1.2E-04 ***** | **pubs_FIRST_RA_JIF_max** |
| ***Mean of "percentile in subject area"*** | **73.5** | **69.7** | **1.6E-03 **** | **pubs_FIRST_RA_pecentile_mean** |
| ***Highest value of "percentile in subject area"*** | **82.0** | **79.2** | **1.9E-02 *** | **pubs_FIRST_RA_pecentile_MAX** |
| *Mean of CNCI* | 2.6 | 2.7 | 8.4E-01 | pubs_FIRST_RA_CNCI_mean |
| *Highest value of "CNCI"* | 3.9 | 4.2 | 5.8E-01 | pubs_FIRST_RA_CNCI_max |
| *Calendar years: EMBL start -> earliest 1st author* | 3.3 | 3.2 | 1.8E-01 | pubs_FIRST_ra_only_publishedfirst |
| *Calendar years: EMBL start -> final 1st author* | 4.8 | 4.8 | 8.9E-01 | pubs_FIRST_ra_only_publishedlast |
| *Mean number of authors* | 6.8 | 7.0 | 5.2E-01 | pubs_FIRST_ra_only_Authors_mean |
| ***Number of co-author research articles*** | **2.0** | **2.4** | **2.8E-03 **** | **coauthor_research** |
| ***Number of publications not classed as research*** | **0.8** | **1.0** | **3.7E-02 *** | **nonresearhc** |

~ statistical significance based on unpaired 2-tailed t-test *<0.05, **<0.01,***<0.001

#### Supplementary table 9: differences in publication statistics by 'industry research' status.

Values that are statistically significant (p<0.05) are highlighted with bold, and those mentioned in paper text are underlined.

| Publication statistics | Became IndRes? no | yes | sig~ | | Field name |
|---|---|---|---|---|---|
| *Number of alumni* | 1811 | 473 | | | |
| For: ***all publications*** *from EMBL work* | | | | | |
| *Number of publications* | 4.6 | 4.5 | 7.2E-01 | | PUBS_ALL_TOTAL |
| *Number of publications with a JIF* | 4.6 | 4.5 | 7.4E-01 | | PUBS_ALL_JIF_count |
| *Mean JIF* | 10.1 | 9.6 | 1.9E-01 | | PUBS_ALL_JIF_mean |
| *Sum of JIFs* | 52.3 | 49.0 | 3.3E-01 | | PUBS_ALL_JIF_SUM |
| ***Highest JIF*** | **20.2** | **18.4** | **2.2E-02** | * | **pubs_ALL_JIF_MAX** |
| *Mean of "percentile in subject area"* | 65.2 | 64.8 | 7.4E-01 | | pubs_All_pecentile_mean |
| *Highest value of "percentile in subject area"* | 86.4 | 86.5 | 9.2E-01 | | pubs_All_percentile_MAX |
| *Mean CNCI* | 2.8 | 2.6 | 3.9E-01 | | PUBS_ALL_CNCI_mean |
| *Highest value of "CNCI"* | 9.1 | 8.8 | 8.2E-01 | | PUBS_ALL_CNCI_max |
| *Calendar years: EMBL start year -> earliest* | 2.4 | 2.4 | 5.6E-01 | | pubs_All_published.first |
| *Calendar years: EMBL start -> final publication* | 5.4 | 5.5 | 9.1E-01 | | pubs_All_published.last |
| *Mean number of authors* | 8.8 | 8.4 | 3.3E-01 | | pubs_All_published_authors_mean |
| For: all **research articles** from EMBL work | | | | | |
| *Number of research articles* | 3.6 | 3.7 | 7.2E-01 | | pubs_RAonly_TOTAL |
| *Number of research articles with a JIF* | 3.6 | 3.7 | 6.9E-01 | | pubs_RAonly_JIF_count |
| *Mean JIF* | 11.1 | 10.4 | 1.2E-01 | | pubs_RAonly_JIF_mean |
| *Sum of JIFs* | 46.7 | 44.8 | 5.3E-01 | | pubs_RAonly_JIF_SUM |
| ***Highest JIF*** | **19.9** | **17.9** | **1.1E-02** | * | **pubs_RA_JIF_max** |
| *Mean of "percentile in subject area"* | 73.7 | 71.8 | 7.0E-02 | | pubs_RAonly_percentile_mean |
| *Highest value of "percentile in subject area"* | 87.7 | 86.6 | 2.6E-01 | | pubs_RA_pecentile_MAX |
| *Mean CNCI* | 3.2 | 2.8 | 1.6E-01 | | pubs_RA_CNCI_mean |
| *Highest value of "CNCI"* | 8.2 | 7.8 | 7.4E-01 | | pubs_RAonly_CNCI_max |
| *Calendar years: EMBL start-> earliest res.article* | 2.7 | 2.6 | 4.3E-01 | | pubs_RAonly_publishedfirst |
| *Calendar years: EMBL start -> final res.article* | 5.5 | 5.5 | 9.1E-01 | | pubs_RAonly_publishedlast |
| *Mean number of authors* | 9.3 | 8.9 | 3.8E-01 | | pubs_RAonly_Authors_average |
| For: **first-author research** articles from EMBL work | | | | | |
| *Number of 1st author research articles* | 1.6 | 1.5 | 2.7E-01 | | pubs_FIRST_ra_only_TOTAL |
| *Number of 1st author research articles with a JIF* | 1.6 | 1.5 | 3.0E-01 | | pubs_FIRST_ra_only_JIF_count |
| *Mean JIF* | 10.7 | 10.1 | 3.1E-01 | | pubs_FIRST_ra_only_JIF_mean |
| ***Sum of JIFs*** | **22.3** | **19.6** | **2.8E-02** | * | **pubs_FIRST_ra_only_JIFsum** |
| ***Highest JIF*** | **14.9** | **13.4** | **4.3E-02** | * | **pubs_FIRST_RA_JIF_max** |
| *Mean of "percentile in subject area"* | 72.7 | 70.7 | 1.5E-01 | | pubs_FIRST_RA_pecentile_mean |
| *Highest value of "percentile in subject area"* | 81.4 | 80.0 | 3.2E-01 | | pubs_FIRST_RA_pecentile_MAX |
| *Mean of CNCI* | 2.7 | 2.4 | 2.1E-01 | | pubs_FIRST_RA_CNCI_mean |
| *Highest value of "CNCI"* | 4.1 | 3.6 | 1.6E-01 | | pubs_FIRST_RA_CNCI_max |
| *Calendar years: EMBL start -> earliest 1st author* | 3.3 | 3.3 | 6.0E-01 | | pubs_FIRST_ra_only_publishedfirst |
| *Calendar years: EMBL start -> final 1st author* | 4.8 | 4.8 | 7.8E-01 | | pubs_FIRST_ra_only_publishedlast |
| *Mean number of authors* | 7.0 | 6.8 | 5.5E-01 | | pubs_FIRST_ra_only_Authors_mean |
| *Number of co-author research articles* | 2.0 | 2.2 | 2.5E-01 | | coauthor_research |
| ***Number of publications not classed as research*** | **0.9** | **0.8** | **3.6E-02** | * | **nonresearhc** |

~ statistical significance based on unpaired 2-tailed t-test *<0.05, **<0.01,***<0.001

**Supplementary table 10: differences in publication statistics by 'non-research science-related' status.**

Values that are statistically significant (p<0.05) are highlighted with bold, and those mentioned in paper text are underlined.

| Publication statistics | Became NonRes? no | yes | sig~ | | Field name |
|---|---|---|---|---|---|
| *Number of alumni* | 1857 | 427 | | | |
| For: ***all publications*** *from EMBL work* | | | | | |
| ***Number of publications*** | **4.8** | **3.2** | **<2e-16** | *** | **PUBS_ALL_TOTAL** |
| ***Number of publications with a JIF*** | **4.8** | **3.2** | **<2e-16** | *** | **PUBS_ALL_JIF_count** |
| *Mean JIF* | 10.1 | 9.6 | 2.0E-01 | | PUBS_ALL_JIF_mean |
| ***Sum of JIFs*** | **55.2** | **34.9** | **4.3E-15** | *** | **PUBS_ALL_JIF_SUM** |
| ***Highest JIF*** | **20.4** | **16.9** | **1.8E-05** | *** | **pubs_ALL_JIF_MAX** |
| *Mean of "percentile in subject area"* | 65.1 | 65.3 | 9.1E-01 | | pubs_All_pecentile_mean |
| ***Highest value of "percentile in subject area"*** | **87.1** | **83.0** | **9.1E-04** | *** | **pubs_All_percentile_MAX** |
| ***Mean CNCI*** | **2.9** | **2.3** | **4.5E-03** | ** | **PUBS_ALL_CNCI_mean** |
| ***Highest value of "CNCI"*** | **9.7** | **5.9** | **2.9E-03** | ** | **PUBS_ALL_CNCI_max** |
| ***Calendar years: EMBL start year -> earliest*** | **2.3** | **2.7** | **5.3E-04** | *** | **pubs_All_published.first** |
| *Calendar years: EMBL start -> final publication* | 5.5 | 5.3 | 3.3E-01 | | pubs_All_published.last |
| *Mean number of authors* | 8.9 | 8.0 | 5.5E-02 | | pubs_All_published_authors_mean |
| For: all research articles from EMBL work | | | | | |
| ***Number of research articles*** | **3.9** | **2.6** | **4.1E-16** | *** | **pubs_RAonly_TOTAL** |
| ***Number of research articles with a JIF*** | **3.9** | **2.6** | **6.4E-16** | *** | **pubs_RAonly_JIF_count** |
| *Mean JIF* | 11.1 | 10.3 | 7.9E-02 | | pubs_RAonly_JIF_mean |
| ***Sum of JIFs*** | **49.5** | **31.4** | **2.0E-14** | *** | **pubs_RAonly_JIF_SUM** |
| ***Highest JIF*** | **20.1** | **16.5** | **1.9E-05** | *** | **pubs_RA_JIF_max** |
| *Mean of "percentile in subject area"* | 73.6 | 71.8 | 1.1E-01 | | pubs_RAonly_percentile_mean |
| ***Highest value of "percentile in subject area"*** | **88.4** | **83.4** | **2.8E-05** | *** | **pubs_RA_pecentile_MAX** |
| ***Mean CNCI*** | **3.2** | **2.3** | **4.3E-06** | *** | **pubs_RA_CNCI_mean** |
| ***Highest value of "CNCI"*** | **8.9** | **4.4** | **3.7E-09** | *** | **pubs_RAonly_CNCI_max** |
| ***Calendar years: EMBL start-> earliest res.article*** | **2.6** | **3.0** | **4.2E-04** | *** | **pubs_RAonly_publishedfirst** |
| *Calendar years: EMBL start -> final res.article* | 5.5 | 5.4 | 5.5E-01 | | pubs_RAonly_publishedlast |
| ***Mean number of authors*** | **9.4** | **8.2** | **8.8E-03** | ** | **pubs_RAonly_Authors_average** |
| For: first-author research articles from EMBL work | | | | | |
| ***Number of 1st author research articles*** | **1.7** | **1.1** | **2.9E-16** | *** | **pubs_FIRST_ra_only_TOTAL** |
| ***Number of 1st author research articles with a JIF*** | **1.7** | **1.1** | **4.3E-16** | *** | **pubs_FIRST_ra_only_JIF_count** |
| ***Mean JIF*** | **10.8** | **9.1** | **4.6E-03** | ** | **pubs_FIRST_ra_only_JIF_mean** |
| ***Sum of JIFs*** | **23.0** | **15.0** | **2.7E-13** | *** | **pubs_FIRST_ra_only_JIFsum** |
| ***Highest JIF*** | **15.2** | **11.5** | **2.2E-06** | *** | **pubs_FIRST_RA_JIF_max** |
| *Mean of "percentile in subject area"* | 72.8 | 69.9 | 5.4E-02 | | pubs_FIRST_RA_pecentile_mean |
| ***Highest value of "percentile in subject area"*** | **81.9** | **77.0** | **7.7E-04** | *** | **pubs_FIRST_RA_pecentile_MAX** |
| ***Mean of CNCI*** | **2.7** | **2.0** | **7.4E-04** | *** | **pubs_FIRST_RA_CNCI_mean** |
| ***Highest value of "CNCI"*** | **4.3** | **2.7** | **6.2E-05** | *** | **pubs_FIRST_RA_CNCI_max** |
| ***Calendar years: EMBL start -> earliest 1st author*** | **3.3** | **3.6** | **1.8E-02** | * | **pubs_FIRST_ra_only_publishedfirst** |
| *Calendar years: EMBL start -> final 1st author* | 4.8 | 4.7 | 2.9E-01 | | pubs_FIRST_ra_only_publishedlast |
| ***Mean number of authors*** | **7.2** | **6.1** | **9.6E-05** | *** | **pubs_FIRST_ra_only_Authors_mean** |
| ***Number of co-author research articles*** | **2.2** | **1.5** | **3.9E-09** | *** | **coauthor_research** |
| ***Number of publications not classed as research*** | **1.0** | **0.6** | **8.6E-07** | *** | **nonresearhc** |

~ statistical significance based on unpaired 2-tailed t-test *<0.05, **<0.01,***<0.001

**Supplementary table 11: differences in publication statistics by ‘non-science role’ status.**

Values that are statistically significant ($p<0.05$) are highlighted with bold, and those mentioned in paper text are underlined.

| | *Became NonRes?* | | | | |
|---|---|---|---|---|---|
| *Publication statistics* | no | yes | sig~ | | Field name |
| *Number of alumni* | 2122 | 162 | | | |
| | | | | | For: ***all publications*** from EMBL work |
| ***Number of publications*** | **4.6** | **3.6** | **2.2E-03** | ** | **PUBS_ALL_TOTAL** |
| ***Number of publications with a JIF*** | **4.6** | **3.5** | **2.1E-03** | ** | **PUBS_ALL_JIF_count** |
| *Mean JIF* | 10.1 | 9.2 | 2.9E-01 | | PUBS_ALL_JIF_mean |
| ***Sum of JIFs*** | **52.4** | **39.4** | **8.3E-03** | ** | **PUBS_ALL_JIF_SUM** |
| ***Highest JIF*** | **20.0** | **16.8** | **1.7E-02** | * | **pubs_ALL_JIF_MAX** |
| *Mean of “percentile in subject area”* | 65.3 | 63.0 | 2.9E-01 | | pubs_All_pecentile_mean |
| ***Highest value of “percentile in subject area”*** | **86.7** | **81.8** | **1.7E-02** | * | **pubs_All_percentile_MAX** |
| *Mean CNCI* | 2.8 | 2.4 | 1.4E-01 | | PUBS_ALL_CNCI_mean |
| *Highest value of “CNCI”* | 9.2 | 6.9 | 2.8E-01 | | PUBS_ALL_CNCI_max |
| *Calendar years: EMBL start year -> earliest* | 2.4 | 2.4 | 8.8E-01 | | pubs_All_published.first |
| *Calendar years: EMBL start year -> last* | 5.5 | 5.1 | 5.8E-02 | | pubs_All_published.last |
| *Mean number of authors* | 8.7 | 9.1 | 6.6E-01 | | pubs_All_published_authors_mean |
| For: all research articles from EMBL work | | | | | |
| ***Number of research articles*** | **3.7** | **2.7** | **2.8E-04** | *** | **pubs_RAonly_TOTAL** |
| ***Number of research articles with a JIF*** | **3.7** | **2.7** | **2.8E-04** | *** | **pubs_RAonly_JIF_count** |
| *Mean JIF* | 11.1 | 9.9 | 2.0E-01 | | pubs_RAonly_JIF_mean |
| ***Sum of JIFs*** | **47.1** | **35.0** | **9.6E-03** | ** | **pubs_RAonly_JIF_SUM** |
| ***Highest JIF*** | **19.6** | **16.4** | **1.9E-02** | * | **pubs_RA_JIF_max** |
| *Mean of “percentile in subject area”* | 73.4 | 71.5 | 2.8E-01 | | pubs_RAonly_percentile_mean |
| ***Highest value of “percentile in subject area”*** | **87.8** | **83.8** | **2.7E-02** | * | **pubs_RA_pecentile_MAX** |
| *Mean CNCI* | 3.1 | 2.6 | 1.5E-01 | | pubs_RA_CNCI_mean |
| *Highest value of “CNCI”* | 8.2 | 6.4 | 3.7E-01 | | pubs_RAonly_CNCI_max |
| *Calendar years: EMBL start-> earliest res.article* | 2.7 | 2.8 | 3.1E-01 | | pubs_RAonly_publishedfirst |
| *Calendar years: EMBL start -> final res.article* | 5.5 | 5.2 | 1.5E-01 | | pubs_RAonly_publishedlast |
| *Mean number of authors* | 9.2 | 9.2 | 9.8E-01 | | pubs_RAonly_Authors_average |
| For: first-author research articles from EMBL work | | | | | |
| ***Number of 1st author research articles*** | **1.6** | **1.1** | **1.1E-05** | *** | **pubs_FIRST_ra_only_TOTAL** |
| ***Number of 1st author research articles with a JIF*** | **1.6** | **1.1** | **1.0E-05** | *** | **pubs_FIRST_ra_only_JIF_count** |
| ***Mean JIF*** | **10.7** | **8.9** | **4.9E-02** | * | **pubs_FIRST_ra_only_JIF_mean** |
| ***Sum of JIFs*** | **22.2** | **14.7** | **6.0E-07** | *** | **pubs_FIRST_ra_only_JIFsum** |
| ***Highest JIF*** | **14.8** | **11.2** | **1.8E-03** | ** | **pubs_FIRST_RA_JIF_max** |
| *Mean of “percentile in subject area”* | 72.5 | 69.6 | 2.2E-01 | | pubs_FIRST_RA_pecentile_mean |
| *Highest value of “percentile in subject area”* | 81.4 | 77.0 | 5.7E-02 | | pubs_FIRST_RA_pecentile_MAX |
| ***Mean of CNCI*** | **2.7** | **1.9** | **2.9E-03** | ** | **pubs_FIRST_RA_CNCI_mean** |
| ***Highest value of “CNCI”*** | **4.1** | **2.4** | **5.2E-06** | *** | **pubs_FIRST_RA_CNCI_max** |
| *Calendar years: EMBL start -> earliest 1st author* | 3.3 | 3.5 | 2.9E-01 | | pubs_FIRST_ra_only_publishedfirst |
| *Calendar years: EMBL start -> final 1st author* | 4.8 | 4.5 | 6.4E-02 | | pubs_FIRST_ra_only_publishedlast |
| ***Mean number of authors*** | **7.1** | **6.1** | **3.4E-02** | * | **pubs_FIRST_ra_only_Authors_mean** |
| ***Number of co-author research articles*** | **2.1** | **1.7** | **2.7E-02** | * | **coauthor_research** |
| *Number of publications not classed as research* | 0.9 | 0.8 | 5.4E-01 | | nonresearhc |

~ statistical significance based on unpaired 2-tailed t-test *<0.05, **<0.01,***<0.001

#### Supplementary table 12: differences in publication statistics by cohort.

Values that are statistically significant (p<0.05) are highlighted with bold, and values mentioned in main text are underlined.

| | Cohort | | | | |
|---|---|---|---|---|---|
| *Publication statistics* | 1997 2004 | 2005 2012 | 2013 2020 | sig~ | Field name |
| *Number of alumni* | 625 | 763 | 896 | | |
| For: ***all publications*** *from EMBL work* | | | | | |
| ***Number of publications*** | **4.7** | **4.2** | **4.7** | **2.8E-02** | ***** **PUBS_ALL_TOTAL** |
| ***Number of publications with a JIF*** | **4.7** | **4.2** | **4.7** | **3.2E-02** | ***** **PUBS_ALL_JIF_count** |
| ***Mean JIF*** | **8.6** | **9.9** | **11.2** | **2.0E-10** | ******* **PUBS_ALL_JIF_mean** |
| ***Sum of JIFs*** | **45.4** | **46.3** | **60.3** | **2.4E-06** | ******* **PUBS_ALL_JIF_SUM** |
| ***Highest JIF*** | **18.5** | **18.6** | **21.7** | **2.4E-05** | ******* **pubs_ALL_JIF_MAX** |
| *Mean of "percentile in subject area"* | 64.6 | 65.4 | 65.3 | 7.9E-01 | pubs_All_pecentile_mean |
| ***Highest value of "percentile in subject area"*** | **86.1** | **84.4** | **88.3** | **1.3E-03** | ****** **pubs_All_percentile_MAX** |
| ***Mean CNCI*** | **2.0** | **2.3** | **3.7** | **1.2E-09** | ******* **PUBS_ALL_CNCI_mean** |
| ***Highest value of "CNCI"*** | **6.1** | **6.6** | **13.3** | **6.6E-07** | ******* **PUBS_ALL_CNCI_max** |
| ***Calendar years: EMBL start year -> earliest*** | **2.2** | **2.6** | **2.4** | **1.3E-05** | ******* **pubs_All_published.first** |
| ***Calendar years: EMBL start year -> last*** | **5.1** | **5.7** | **5.5** | **6.5E-05** | ******* **pubs_All_published.last** |
| ***Mean number of authors*** | **5.8** | **8.0** | **11.3** | **<2e-16** | ******* **pubs_All_published_authors_me** |
| For: all **research articles** from EMBL work | | | | | |
| *<u>Number of research articles</u>* | 3.8 | 3.4 | 3.7 | 1.2E-01 | pubs_RAonly_TOTAL |
| *Number of research articles with a JIF* | 3.8 | 3.4 | 3.7 | 1.2E-01 | pubs_RAonly_JIF_count |
| ***Mean JIF*** | **9.4** | **10.6** | **12.4** | **5.6E-11** | ******* **pubs_RAonly_JIF_mean** |
| ***Sum of JIFs*** | **40.0** | **41.4** | **54.9** | **2.3E-07** | ******* **pubs_RAonly_JIF_SUM** |
| ***Highest JIF*** | **17.9** | **18.3** | **21.5** | **3.0E-06** | ******* **pubs_RA_JIF_max** |
| *Mean of "percentile in subject area"* | 72.2 | 73.1 | 74.2 | 1.5E-01 | pubs_RAonly_percentile_mean |
| ***Highest value of "percentile in subject area"*** | **87.0** | **86.2** | **88.9** | **1.5E-02** | ***** **pubs_RA_pecentile_MAX** |
| ***Mean CNCI*** | **2.2** | **2.6** | **4.1** | **8.7E-11** | ******* **pubs_RA_CNCI_mean** |
| ***Highest value of "CNCI"*** | **5.0** | **6.3** | **11.8** | **1.3E-07** | ******* **pubs_RAonly_CNCI_max** |
| ***Calendar years: EMBL start -> earliest*** | **2.3** | **2.9** | **2.7** | **4.6E-09** | ******* **pubs_RAonly_publishedfirst** |
| ***Calendar years: EMBL start year -> last res.*** | **5.1** | **5.8** | **5.4** | **3.2E-06** | ******* **pubs_RAonly_publishedlast** |
| ***Mean number of authors*** | **5.9** | **8.3** | **12.2** | **<2e-16** | ******* **pubs_RAonly_Authors_average** |
| For: **first-author research articles** from EMBL work | | | | | |
| ***<u>Number of 1st author research articles</u>*** | **1.9** | **1.5** | **1.4** | **1.3E-10** | ******* **pubs_FIRST_ra_only_TOTAL** |
| ***No. 1st author research articles with JIF*** | **1.9** | **1.5** | **1.3** | **5.5E-11** | ******* **pubs_FIRST_ra_only_JIF_count** |
| ***Mean JIF*** | **9.2** | **10.0** | **12.1** | **3.0E-07** | ******* **pubs_FIRST_ra_only_JIF_mean** |
| *Sum of JIFs* | 22.7 | 20.3 | 22.3 | 1.6E-01 | pubs_FIRST_ra_only_JIFsum |
| ***Highest JIF*** | **14.1** | **13.7** | **15.7** | **1.7E-02** | ***** **pubs_FIRST_RA_JIF_max** |
| *Mean of "percentile in subject area"* | 71.8 | 72.3 | 72.7 | 8.0E-01 | pubs_FIRST_RA_pecentile_mean |
| *Highest value of "percentile in subject area"* | 82.0 | 80.3 | 81.2 | 4.4E-01 | pubs_FIRST_RA_pecentile_MAX |
| ***Mean of CNCI*** | **2.0** | **2.5** | **3.1** | **9.6E-04** | ******* **pubs_FIRST_RA_CNCI_mean** |
| *Highest value of "CNCI"* | 3.6 | 3.9 | 4.5 | 2.3E-01 | pubs_FIRST_RA_CNCI_max |
| ***Calendar years: EMBL start -> earliest 1st*** | **2.8** | **3.5** | **3.5** | **4.9E-12** | ******* **pubs_FIRST_ra_only_publishedfi** |
| ***Calendar years: EMBL start -> last 1st author*** | **4.5** | **5.0** | **4.9** | **7.0E-05** | ******* **pubs_FIRST_ra_only_publishedla** |
| ***Mean number of authors*** | **4.8** | **6.1** | **9.4** | **<2e-16** | ******* **pubs_FIRST_ra_only_Authors_m** |
| ***Number of co-author research articles*** | **1.9** | **1.9** | **2.4** | **5.4E-04** | ******* **coauthor_research** |
| ***Number of publications not classed as*** | **0.9** | **0.8** | **1.0** | **5.5E-03** | ****** **nonresearhc** |

~ statistical significance based on ANOVA analysis *<0.05, **<0.01,***<0.001

**Supplementary Methods**

**Identification of alumni for inclusion in the study:**

- Postdocs who spent at least 12-months at EMBL were identified through our alumni records, HR and postdoc office records.
- PhD students who graduated from EMBL's international PhD programme were identified through our alumni, graduate office and HR records. The thesis defence year was identified using graduate office records.
- Deceased alumni were omitted from the study.

**Basic information**

- Information about gender, nationality and time spent at EMBL was collated from information held in HR records, alumni and graduate and postdoc office records. The data was imported and collated in a password-protected custom-made filemaker database. Information that could be used to easily identify the alumnus were pseudonymized after data compilation (e.g. name, unit & group at EMBL, nationality).
- For PhD students the start year was the start of their PhD and end year was the thesis defence year.
- For postdocs the start and end years were based on the start and end year of their EMBL postdoc contract, including any extensions to that postdoc contract.
- Additional positions at EMBL (e.g. a bridging postdoc contract for former predocs, or a later staff position) were included in the career tracking as a subsequent position, not as a continuation of their time as a PhD student or original postdoc.

**Data collection – career information**

- Career information was identified by carrying out a search for publicly available information (LinkedIN profile, researchgate profile, ORCID record and biosketch or CV on employer or personal website, as well as inclusion on publicly available staff lists). The search engine google was used with the search terms
    - [name]
    - [name] + EMBL OR "European molecular biology laboratory"
    - [name] + last known employer
        - the last known employer was taken from information provided to EMBL's alumni office. This information was, however, not included unless it could be confirmed with other publicly available information.
- Identity was confirmed only if the publicly available information mentioned having worked/studied at EMBL or if a publicly available listing of their publications was able to confirm the matching identity.
- Basic details of the positions provided in online career information (country, start year and end year) were added to the filemaker database along with a classification of the position (see classification of positions). A calculated field allowed retrieval of the role classification at specific timepoints after EMBL. Where two positions were held in the same calendar year, the most recent position was retrieved.
- Where this was available, we also noted for each postdoc alumni, the year the PhD was awarded.
- If the current position found on personal webpages or profiles appeared to be a temporary academic position (e.g. postdoc or similar) and they had held the position since before 2019, we confirmed this was up-to-date by cross-checking institutional websites. If group pages do not list lab members, we instead checked that there was a publication within the last calendar year with this affiliation.
- Data on positions was initially collected in March 2017-December 2017 for fellows leaving in 2016 or earlier; and updated in May- September 2021 for all fellows. Country names were pseudonymized after data curation.

**Classification of positions**

- A detailed three-level classification of roles was carried out based on a published taxonomy https://f1000research.com/articles/9-8/v2/. The three-levels of classification include "job sector", "job type" and "job function". As it was not always possible to confirm whether academic positions in Europe were research or teaching focussed, whether specific job titles in industry were at the group leader level, and whether roles in industry R&D were computational biology positions, some adaptations were made. These are detailed in the footnotes of supplementary table 2-3.
- A summary classification into one of six main types of role (independent group leader (AcPI); academic postdoc (AcPD); other academic position (AcOt); industry research (IndR); science related (SciR); and non-science related (NonSci)) was then calculated based on the detailed classification.
    - AcPI includes those with the job function "Independent research group leader"
    - AcOt includes with the job function "Other academic position"

- AcPD includes those in the sector academia, with the job function “postdoctoral”
- IndR includes alumni:
  - with the job function “R&D scientist”
  - with the job function “postdoctoral” and sector “Industry / for-profit”
  - with the career type “research / teaching”, sector “Industry / for-profit”, and function "Business development, consulting, and strategic alliances" OR “Entrepreneurship” (this captures senior leadership roles overseeing industry research strategy, and founders of research- focused start-ups, including small contract research organisations).
- NonRes includes
  - alumni with job type="Science related non-research”
  - alumni with job type “Research / teaching” who were *not* employed in academia or had a job function that linked them to industry research [see IndR] (this captures those who are working in teaching outside academia (e.g. secondary education))
- NonSci includes all remaining positions

**Data collection – publication information**

- Data on all publications linked to EMBL were exported from the Web of Science incites database on 30 June 2021 - including the publication year, research area, type of publication, citation metrics such as percentile in subject area, and journal impact factor for each publication.
  - Data from the main web of science database (WOS) for all EMBL publications was also exported on 30 June 2021, and the full author list and affiliations added by matching the web of science ascension numbers.
  - A small number of publications were listed in WOS but not incites – the article type from WOS was manually added in these cases.
- Alumni in this study were matched to the publications identified from Web of Science as follows:
  - publications during the years ECRs were at EMBL were assigned to individuals without further validation if:
    - the paper was already registered to them in EMBL’s internal publication database
    - there was a match between an author and the individual’s last name and initial, or known variants of their name (e.g. previous surname for alumni who changed their name on marriage) – AND their EMBL supervisor’s name was also on the paper
  - additional publications with matching name & first initial that were published during or after the ECR’s EMBL contract were manually checked and assigned to ECRs if the publication data (full name, affiliation etc) supported that this was a publication from that person and stemmed from EMBL work.
    - an exception was made if the publication was more than 2-calendar years after EMBL and the person was known to have remained or returned to EMBL in a research-based position (e.g. staff scientist) at that time-point.
  - for ECRs with potential name variations (e.g. double last name or special characters), publications were also added manually where appropriate.
- Joint first authorship was established by manually checking all publications where an ECR was the second author. Joint first-authorship contributed to 14.6% of articles classed as first-author. In a small proportion of cases the ECR was the senior (last) author on a research article, and we included this as equivalent to a first-author article [4.8% of articles classed as first-author].
- Once data curation was complete, the publication records were pseudonymized by replacing the web of science ID with a pseudonym, and deletion of additional identifying information not required for the final analysis (e.g. author lists, affiliations, journal name, article title).

**Classification of supervisor**

- The supervisor was classed as senior if, in the year the ECR left, their supervisor was a senior scientist OR had spent >10-years at EMBL.
